# FlexiTAC enables controllable PROTAC linker generation across diverse structural settings using a Bayesian flow network with posterior guidance

**DOI:** 10.64898/2026.08.26.747172

**Authors:** Yuzhang Li, Yupu Zhao, Lixin Zhou, Changyin Huang, Qisheng Xu, Yang Chen, Zheng Qin, Kelong Fan, Jing Yang, Duanhua Cao

## Abstract

Linker chemistry and conformation are central determinants of PROTAC activity, shaping ternary-complex geometry, cooperativity, target-lysine presentation and cellular permeability. Existing linker generators often lack explicit control over linker flexibility, require predefined attachment sites and linker lengths, or produce structures that demand substantial geometric correction, limiting their utility in practical PROTAC design. Here we introduce FlexiTAC, a Bayesian flow network that jointly generates linker atom types and coordinates from the warhead and E3-ligase-ligand contexts. We also assemble PROTAC-3D, a quality-controlled collection of 63,554 component-resolved PROTAC structures for model training, and PROTAC-Bench, which covers molecular quality, fragment preservation, geometric fidelity, conformational stability, fragment awareness, rediscovery and sampling efficiency. Compared to the best 3D baseline models, FlexiTAC improves validity by 12.0-12.7% and achieves the highest PoseBusters pass rate of 79.5%-80.0%. A differentiable guidance module shifted generated linkers along a conformational ensemble-derived rigidity axis without retraining the generator. In silico case studies further show that the model can accept crystal-derived, redocked or predicted structural inputs. Together, FlexiTAC, PROTAC-3D and PROTAC-Bench establish an integrated and reproducible framework for data-driven PROTAC linker design, combining controllable structure-conditioned generation with standardized training data and evaluation protocols. This framework expands the linker chemical and conformational space accessible to computational exploration, provides a foundation for future method development and enables the systematic generation of structure-conditioned linker designs with tunable conformational flexibility.

## Introduction

Artificial intelligence (AI) is reshaping drug discovery by turning molecular design into a more data-driven and structure-aware process^1–5^. This shift is especially important for structure-based drug design (SBDD), where activity is governed not only by chemical composition, but also by three-dimensional (3D) shape, conformational preference and molecular placement in a biological environment. Generative drug design increasingly requires models to coordinate discrete chemical identity with continuous 3D geometry while respecting fixed molecular context. PROTAC linker design makes this challenge explicit: a linker must connect two preselected ligands without disrupting their poses, while satisfying chemical and conformational constraints.

Proteolysis-targeting chimeras (PROTACs) induce proximity between an E3 ubiquitin ligase and a protein of interest (POI), leading to ubiquitination and proteasomal degradation^6–8^. This event-driven mechanism endows PROTACs with three key advantages: overcoming the limitations of traditionally “undruggable” targets, circumventing target protein resistance, and potentially achieving therapeutic effects at lower doses^9^. The field reached a clinical milestone with the approval of vepdegestrant^10^. As targeted protein degradation (TPD) has grown, AI methods have also begun to support PROTAC discovery. DeepPROTACs^11^ and PROTAC-STAN^12^ predict degradation activity, DeepTernary^13^ models ternary complexes, Chemistry-42^14^ explores generative degrader design, and DeepDegradome^15^ supports fragment-based generation. Together, these studies illustrate the expansion of computational support across the PROTAC discovery workflow.

Together with attachment sites, a PROTAC linker constrains ligand distance, relative orientation and conformational freedom, while also affecting size, polarity, permeability and metabolic stability. Linker design is therefore a coupled chemical and structural optimization problem, and no single rigidity or geometry is expected to be optimal across targets. Several deep-learning approaches have been developed for PROTAC linker design. Early approaches, such as DeLinker^16^, defined the task as a 2D graph-based generation task. Reinforcement learning strategies were introduced in frameworks like PROTAC-RL^17^. More recently, generative diffusion algorithms, exemplified by DiffLinker^18^ and LinkerNet^19^, move the problem into the SBDD paradigm and leverage a diffusion algorithm to learn the data distribution of linker structures. Despite substantial progress, current methods still face several limitations. Generated molecules may fail chemical-validity, fragment-preservation or geometric-quality checks, provide insufficient coverage of bioactive chemical space and offer limited control over linker properties. We attribute the current limitations in model development to three main challenges. First, data scarcity further limits model development. Curated resources such as PROTAC-DB^20–22^ provide valuable activity and molecular information, but their scale is limited for training structure-based generative models. Larger patent-derived repositories, including PROTAC-PatentDB^23^, contain more PROTAC entries, but they are not consistently segmented into warhead, linker and E3-ligand components, and lack standardized conformations. Second, existing benchmarks and metrics do not adequately reflect practical linker-design capability. They focus mainly on basic molecular validity and diversity, while insufficiently evaluating fragment preservation, 3D geometry, generalization, sampling efficiency, condition awareness and controllability, thereby providing limited guidance for model development. Consequently, strong performance on existing metrics does not necessarily indicate that a model can generate chemically faithful, geometrically plausible and practically useful linkers under realistic design conditions. Third, controllability is a separate challenge. Flexible linkers can broaden conformational sampling, whereas conformational constraint can reduce the accessible ensemble and alter ternary-complex organization^24^. However, rigidity can also change polarity, permeability and synthetic accessibility, and its effect on degradation is system dependent. A useful generator should therefore tune linker mechanics while exposing these correlated property changes.

To address these challenges, we constructed PROTAC-3D, to our knowledge the largest conformational dataset of PROTACs, with rigorous component decomposition and standardized training and test splits. We further established PROTAC-Bench with new test sets and evaluation metrics to better reflect performance in practical linker-design scenarios. Finally, we developed FlexiTAC, a 3D linker-generation model that enables efficient generation and explicit control over linker flexibility.

PROTAC-3D contains more than 60,000 standardized, decomposed and conformationally processed examples from literature- and patent-derived records. Because many source entries contain only full-molecule SMILES, we developed a multi-stage decomposition, quality-control and repair pipeline to recover warhead-linker-E3-ligand annotations. This process increased the number of usable fragment-conditioned examples by approximately tenfold relative to PROTAC-DB 3.0 alone, substantially expanding the degrader chemotypes represented during training. To further improve 3D generation and chemical-space coverage, we derived approximately one million PROTAC-sized fragment-linking examples from ChEMBL3D for pretraining. This enabled the model to learn general chemical and geometric principles of fragment connection before PROTAC-specific fine-tuning. PROTAC-Bench complements these training resources with standardized fragment-pair-level test settings and a hierarchical evaluation protocol across diverse design scenarios. Beyond basic properties such as validity, uniqueness and novelty, PROTAC-Bench evaluates whether a model preserves the conditioning fragments, generalizes across unseen warheads, E3 ligands and fragment pairs, and adapts to different anchor separations and linker lengths. Its metrics further distinguish graph-level correctness from 3D usability by assessing reference recovery and coverage, linker-motif diversity, geometric fidelity and conformational stability under force-field relaxation. In addition, test-time scaling analyses use sampling-budget curves to quantify how efficiently a model recovers reference-like chemotypes. Together, these evaluations provide a more stringent and practically relevant assessment of conditional linker generation than chemical-validity metrics or isolated case studies alone.

Finally, building on these data and benchmarking resources, we developed FlexiTAC, a fragment-conditioned model for PROTAC linker generation. FlexiTAC uses a Bayesian Flow Network (BFN)^25–28^ to jointly model discrete atom types and continuous 3D coordinates. Given two molecular fragments (a POI-binding warhead and an E3-ligase ligand), the model generates the missing linker while preserving the fragment context. This formulation provides a natural way to model the mixed discrete-continuous nature of molecular structure and enables fast sampling. To enable explicit control over linker flexibility, we developed a conformational ensemble-derived rigidity score (CERScore) that integrates three complementary determinants of linker flexibility: local topological freedom quantified by rotatable bonds, end-to-end stability measured by fluctuations in anchor distance, and global shape stability measured by long-range intralink distance fluctuations across generated conformers. Unlike single-conformer or topology-only descriptors, CERScore captures how consistently a linker maintains its overall geometry within an accessible conformational ensemble. By training a differentiable module to predict CERScore, we further enable posterior gradient guidance^26,29^ for linker flexibility control. During sampling, gradients from the CERScore predictor steer the Bayesian parameters toward more flexible or more rigid linkers, allowing users to tune generated structures for early exploration or late-stage optimization.

We compared FlexiTAC with seven representative baselines spanning two-dimensional graph generation, 3D linker generation, equivariant diffusion and transferable molecular generation. These models were retrained on the same PROTAC-3D training split using their original objectives; methods available only as pretrained models were evaluated using their released checkpoints.

Among the evaluated 3D generators, FlexiTAC achieved the strongest overall balance of chemical validity, fragment preservation and conformer similarity, and maintained robust generation across anchor separations ranging from 3.1 Å to 27.1 Å. It achieved PoseBusters pass rates of 79.5-80.0% and more closely reproduced reference bond-length, bond-angle and torsion-angle distributions. After force-field relaxation, the corresponding proportions for FlexiTAC (ChEMBL) were 28.7%, 24.1% and 23.4% across the three test sets, respectively. FlexiTAC also recovered a broader fraction of known PROTAC chemotypes and achieved higher reference coverage within limited sampling budgets. Posterior-gradient guidance produced graded shifts along the conformer-derived rigidity axis, accompanied by the expected changes in rotatable bonds and ring content, while maintaining chemically plausible molecular-property distributions. We further evaluated FlexiTAC in case studies using fragment poses derived from ternary crystal structures, molecular redocking and HDOCK-predicted protein complexes, together with an atypical macrocycle-like PROTAC topology. The model generated chemically connected and geometrically plausible linkers across these settings, supporting its adaptability to structural inputs of different provenance and its potential integration into structure-enabled PROTAC design workflows.

To separate architectural gains from data-scale effects, we retrained FlexiTAC on the legacy ZINC and PROTAC-DB 1.0 benchmarks and compared it with released baseline checkpoints under established evaluation protocols. Its competitive performance and improvements in recovery and conformational quality support a contribution from the Bayesian-flow architecture beyond the expanded PROTAC-3D dataset.

Together, our work establishes an integrated framework for systematic and controllable PROTAC linker design. FlexiTAC provides, to our knowledge, the first 3D linker-generation model with explicit control over conformational flexibility. By translating the spatial arrangement of POI-binding and E3-ligase-binding ligands into chemically and conformationally plausible linker hypotheses, FlexiTAC connects structural information with medicinal-chemistry exploration, while its rigidity guidance enables deliberate sampling across flexible and conformationally constrained designs. Beyond the model itself, PROTAC-3D provides standardized, component-resolved training data spanning diverse degrader chemotypes, whereas PROTAC-Bench enables hierarchical evaluation of chemical validity, geometric plausibility, generalization, sampling efficiency and controllability. These resources provide a reproducible foundation for developing, comparing and diagnosing future linker-generation methods. By aligning model development with task-specific data and application-relevant evaluation, this model-data-benchmark paradigm may accelerate the iterative optimization and facilitate the exploration of new target-E3-ligase combinations, particularly when structural information is limited or derived computationally. Although developed for PROTACs, the underlying formulation of mixed discrete-continuous, structure-constrained molecular completion may extend to other heterobifunctional modalities and fragment-linking problems, providing a bridge between structural biology, generative modeling and medicinal chemistry. Ultimately, such advances may broaden the range of proteins and biological mechanisms accessible to targeted degradation and support the development of new therapeutic strategies.

## Results

### PROTAC-3D and PROTAC-Bench enable large-scale training and rigorous evaluation of linker-generation models

Reliable fragment boundaries were essential because segmentation errors would otherwise be learned as incorrect molecular connection rules. We therefore evaluated all candidate segmentations using a unified quality-control workflow (**Fig.1a**). This workflow jointly assessed molecular validity, fragment size, anchor atoms labelling, linker connectivity, linker ring count and obvious residual linker-like chains in the terminal fragments. Segmentations that failed these checks were labelled as quality-failed and passed to a repair workflow. This step retained chemically recoverable records while allowing unreliable boundaries to be excluded later. The repair workflow combined reference-based re-segmentation with iterative model refinement. Warhead and E3-ligand vocabularies initialized from PROTAC-DB 3.0 were expanded with terminal fragments from quality-passed segmentations and used to identify boundary bonds in complete PROTACs. Molecules with unambiguous matches in the database were re-cut and re-evaluated using the same quality-control rules. The resulting high-quality segmentations were then used to fine-tune the Transformer splitter, which, together with the XGBoost fallback, was reapplied to the remaining problematic records. The final dataset was generated by merging all segmentations that passed the quality checks in either round. When multiple valid segmentations were obtained for the same molecule, we selected the segmentation with the largest linker heavy-atom count. In total, PROTAC-3D comprised 63,554 entries, encompassing 10,649 unique warheads, 5,564 unique E3 ligands and 28,587 unique warhead-E3-ligand pairs after SMILES canonicalization. This represented an approximately tenfold increase over PROTAC-DB 3.0 alone. This enabled the downstream model to learn from a larger and more chemically diverse design space.

To enable the model to learn linker-connection patterns from physically plausible, low-energy molecular geometries, we generated conformers for each retained full-length PROTAC using RDKit^30^. Explicit hydrogen atoms were added before conformation generation, and 20 candidate conformers were generated using the EmbedMultipleConfs module in RDKit. Each conformer was optimized with the MMFF94 force field in RDKit^30^ for up to 1,000 iterations. Among conformers that converged, we retained the lowest-energy structure and removed explicit hydrogen atoms. The curated warhead, linker and E3 ligand SMILES were then matched back to the optimized molecule to assign atom-level fragment identities. We retained only molecules for which all three components were present, non-overlapping and together covered every heavy atom. The resulting data containing coordinates, atom identities, fragment masks and anchor annotations are used for model training and evaluation.

Dataset scale alone does not establish whether the resulting collection occupies chemical space relevant to PROTAC design. We therefore compared PROTAC-3D with PROTAC-DB 3.0 across structural and physicochemical descriptors (**Supplementary Figs. 1-4**). The two datasets occupied broadly overlapping physicochemical space, whereas PROTAC-3D provided substantially greater structural coverage and contained more cyclic linker motifs. This expanded representation exposed the model to a broader range of linker flexibility and conformational constraints. Although PROTAC-3D substantially expands the available training data, PROTAC-specific molecules remain scarce relative to general chemical corpora, and the conformational fidelity of their generated structures remains limited. To this end, we constructed a large corpus of PROTAC-sized quasi-PROTAC molecules from ChEMBL3D, which provides molecular conformations optimized using high-accuracy quantum-chemical calculations. Following the fragmentation strategy introduced in DeLinker, each molecule was divided into two terminal fragments and an intervening linker. This procedure yielded 1,000,450 pretraining examples, enabling the model to learn chemically diverse fragment-linker compositions and physically plausible geometric connection patterns before PROTAC-specific fine-tuning.

Meaningful comparison between linker-generation models is difficult when studies use different data splits, sampling budgets and metric definitions. We therefore developed PROTAC-Bench, a unified benchmark for sequence-based and structure-based PROTAC linker-generation models (**Fig. 1b**). To distinguish component-level generalization from fragment-pair memorization, terminal fragments were canonicalized and PROTACs sharing the same warhead-E3 ligand pair were assigned to a common system. Systems were then classified according to the presence of their terminal fragments in the training set. Both-unseen systems contained neither terminal fragment in the training data, E3-unseen systems combined a previously observed warhead with an unseen E3 ligand, and warhead-unseen systems represented the converse setting.

**Figure 1.**
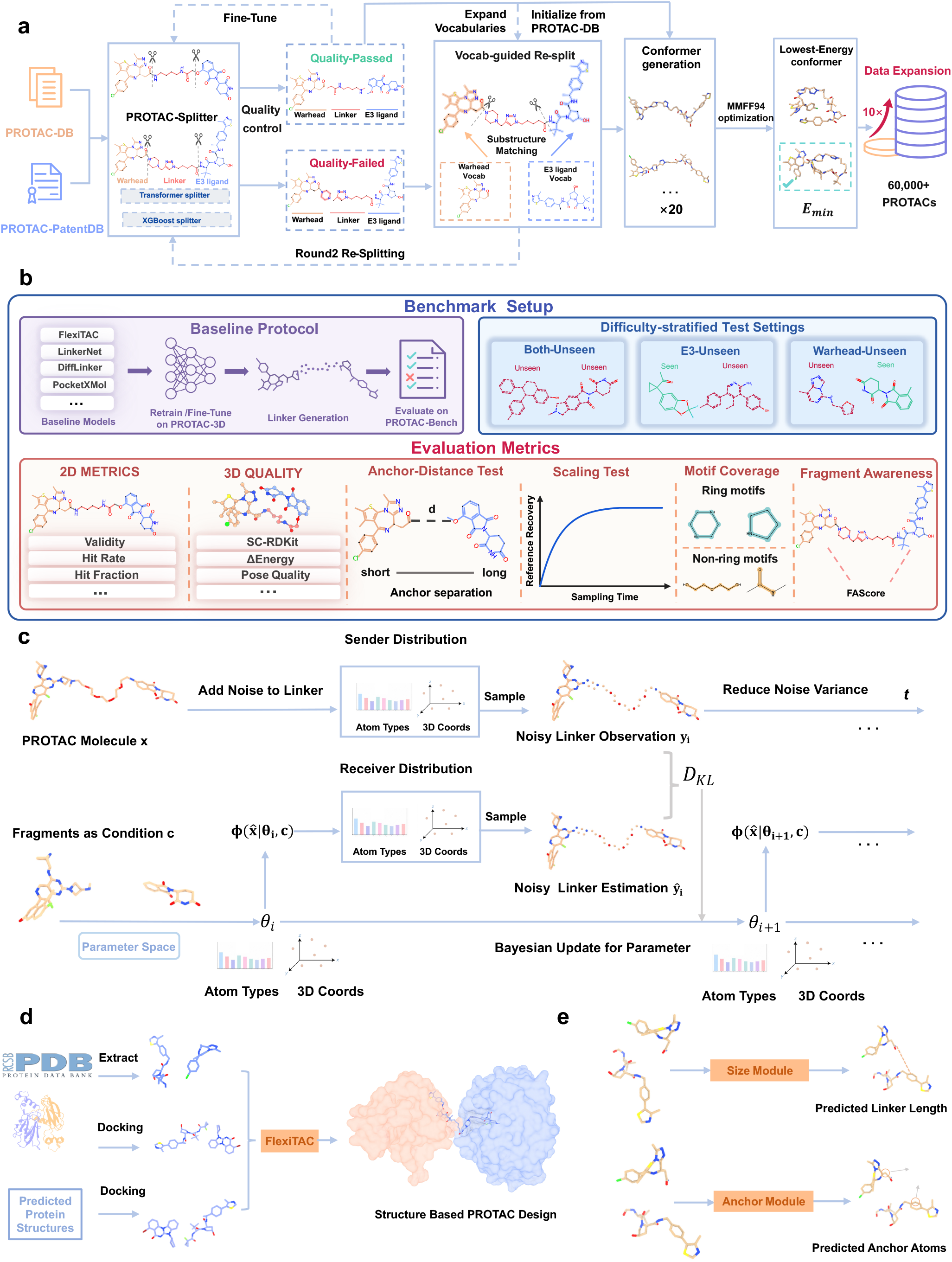
| Construction of PROTAC-3D, the PROTAC-Bench evaluation framework and the FlexiTAC architecture. **a,** Construction of PROTAC-3D from PROTAC-DB and PROTAC-PatentDB. Full PROTACs are separated into warhead, linker and E3-ligand components, followed by quality control and iterative re-splitting. Vocabulary-guided substructure matching expands the chemical coverage of fragment definitions. 3D conformers are then generated, optimized using MMFF94 and ranked by energy to retain the lowest-energy conformer. **b,** Overview of the PROTAC-Bench testing and evaluation framework. FlexiTAC and other baseline models were retrained or fine-tuned on PROTAC-3D and assessed under a unified protocol. Generalization was evaluated across both-unseen, E3-unseen and warhead-unseen settings. PROTAC-Bench combines 2D and 3D assessments with anchor-distance, scaling and motif-coverage tests to evaluate chemical validity, structural quality, robustness, sampling efficiency, linker diversity and Fragment-awareness. **c,** Overall architecture of FlexiTAC **d,** Structural inputs supported by FlexiTAC. Fragment poses can be extracted from experimentally resolved ternary complexes, obtained by docking into experimental protein structures, or derived by docking into predicted protein-protein complexes. **e,** Optional linker-size and anchor-prediction modules. The modules estimate the required linker length and rank candidate attachment atoms when these annotations are unavailable.

To support reliable recovery analysis, we prioritized fragment-pair systems containing larger numbers of unique reference linkers. The resulting benchmark comprised 3,516 reference PROTACs across 130 systems: 555 in 30 both-unseen systems, 1,083 in 50 E3-unseen systems and 1,878 in 50 warhead-unseen systems.

All methods were evaluated under consistent sampling conditions using complementary 2D and 3D metrics spanning chemical validity, reference recovery, conformational quality and physical plausibility. PROTAC-Bench further included anchor-distance, scaling and motif-coverage tests to probe geometric robustness, sampling efficiency and structural diversity beyond exact reference matching. In addition, PROTAC-Bench includes the Fragment-Aware Score (FAScore) to quantify a linker-design model’s awareness of fragment condition. Together, these evaluations assessed model generalization across fragment novelty and distinct linker-design challenges. Full construction procedures, sampling protocols and metric definitions are provided in Methods, “PROTAC-Bench construction and evaluation”.

### FlexiTAC architecture for flexible-input and controllable PROTAC linker generation

FlexiTAC is a fragment-conditioned generative model for PROTAC linker design. It receives an E3-ligase ligand and a POI-binding warhead with fixed atom types, coordinates and attachment information. The model predicts the atom identities and 3D coordinates of a linker that reconnects the two fragments. This formulation reflects practical PROTAC design, in which the fragments define the biological and geometric constraints while the linker unifies them into one molecule.

The core generator is based on a Bayesian Flow Network (BFN) (**Fig. 1c**). The BFN represents discrete atom identities and continuous coordinates through time-dependent distribution parameters. These parameters jointly encode categorical probabilities for atom types and probability distributions for coordinates. During training, the true linker is converted into noisy observations at randomly sampled time points. Early observations contain little information, whereas later observations increasingly constrain the molecular structure according to a time-dependent noise schedule. At each training step, a sender distribution perturbs the true linker at the sampled noise level. The current linker parameters and fixed fragment context are then processed by an SE(3)-equivariant graph neural network. The network predicts a linker estimate that defines a receiver distribution at the same noise level. Minimizing the divergence between the sender and receiver distributions teaches the model to update linker chemistry and geometry while preserving the fragment context. Fragment and linker atoms are represented within a shared molecular graph. Equivariant message passing allows the evolving linker to respond to the positions and chemical environments of both fragments and their attachment atoms. Sampling begins with uniform atom-type probabilities and a Gaussian coordinate prior. The model then applies a sequence of Bayesian updates that progressively reduce uncertainty. At each step, FlexiTAC’s network decodes the current parameter state and predicts the information required for the next update. After the final step, atom identities and coordinates are decoded, chemical bonds are reconstructed, and the linker is joined to the unchanged input fragments.

FlexiTAC is designed to accommodate fragment poses obtained from structural pipelines of different origins (**Fig. 1d**). In information-rich settings, fragment poses can be extracted from an experimentally resolved ternary complex or from the conformation of a known PROTAC. When these structures are unavailable, fragment poses can instead be obtained by redocking ligands into experimental protein structures. They can also be derived by docking fragments into computationally predicted POI-E3 complexes. This design enables the same generator to operate across crystal-derived, redocked and predicted structural inputs. In our case studies, FlexiTAC generated connected PROTACs while preserving the prescribed fragment geometries. The resulting ternary-complex models also showed favorable predicted energy estimates.

To support diverse application scenarios, we equipped FlexiTAC with several auxiliary modules that accommodate structural inputs of varying provenance and completeness. Linker-size and anchor predictors infer the required linker length and candidate attachment sites when these annotations are unavailable, whereas posterior-gradient guidance enables tunable control over linker flexibility during inference. Together, these modules allow FlexiTAC to operate across fully specified and partially annotated design settings with different conformational objectives (**Fig. 1e**). Architectural, training details and evaluation results for these auxiliary modules are provided in the Methods a nd **Supplementary Fig. 5**.

### FlexiTAC designs Chemically Reliable Linker Across Diverse Fragment Geometries

We first evaluated whether FlexiTAC could generate chemically valid molecules with physically plausible 3D conformations. We evaluated two versions of FlexiTAC on PROTAC-Bench,one trained directly on PROTAC-3D: FlexiTAC (scratch), and another first trained on the quasi-PROTAC pre-training data and then fine-tuned on PROTAC-3D: FlexiTAC (ChEMBL), and concurrently trained baseline models using the PROTAC-3D data for a fair comparison. The molecular-property results are summarized in **Fig. 2a,b**. Across the general molecular-property metrics on PROTAC-Bench, the two versions of FlexiTAC reached validity rates of 87.2% and 87.9% respectively, the highest among 3D generative models and close to the best 2D baselines and retained strong functional-group and ring-type consistency and achieved the highest Shape and Color similarity score(SC-RDKit)>0.95 pass rates, surpassing DiffLinker by 5.8 and 7.5 percentage points, respectively. The atom-stability analysis gave a similar picture, with FlexiTAC (scratch) and FlexiTAC (ChEMBL) reaching 96.8% and 97.5%, respectively, above LinkerNet, PocketXMol and 3DLinker.

**Figure 2.**
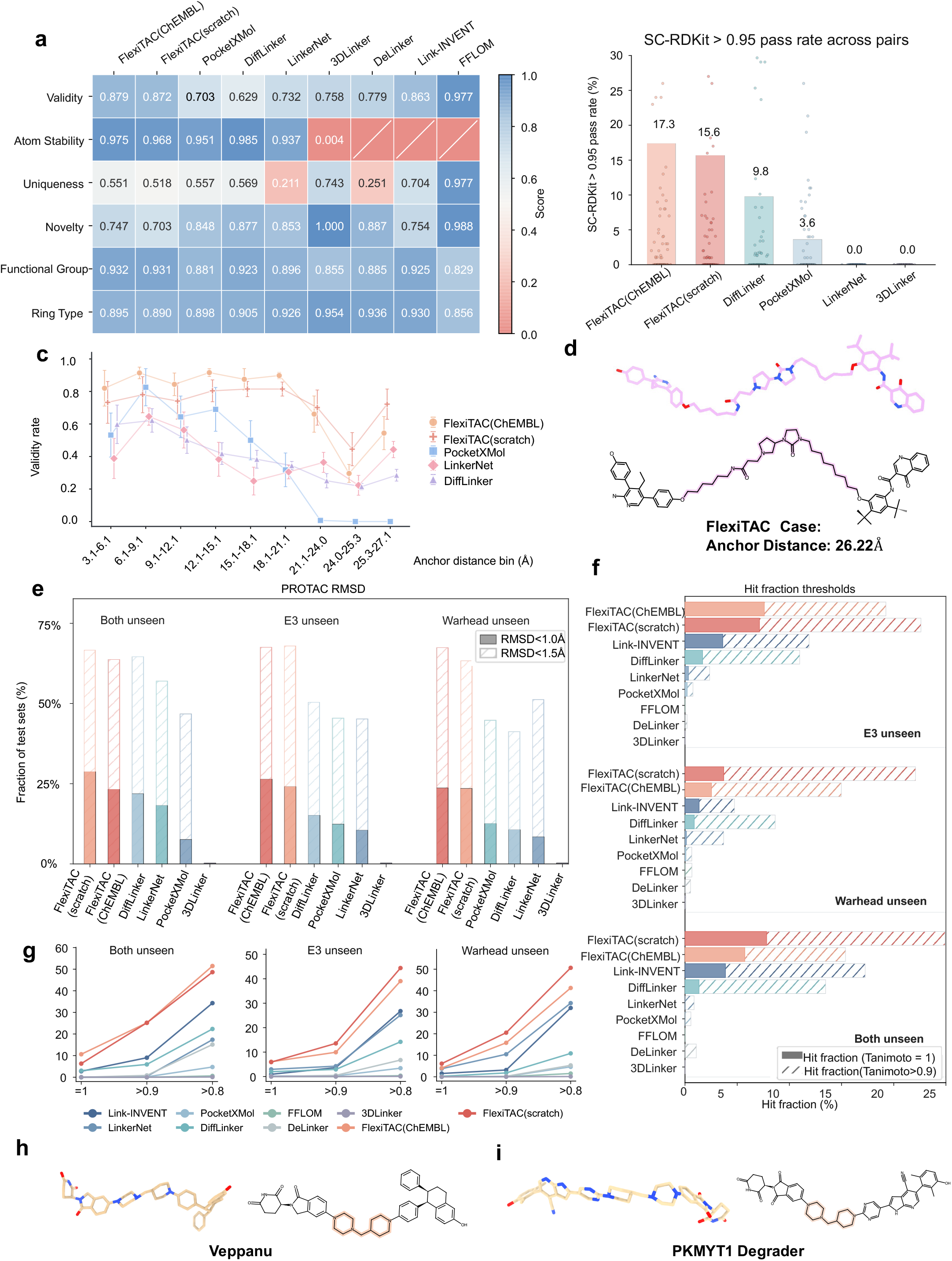
| Molecular quality, linker distance test performance and rediscovery ability of FlexiTAC. **a,** General molecular property comparison across baseline models. Atom stability, uniqueness, novelty, functional-group coverage and ring-type coverage, indicating balanced chemical quality rather than improvement in only one metric. **b,** SC-RDKit pass rate across generated samples. SC-RDKit metric evaluates the geometric and chemical similarity between the ground-truth and generated PROTACs. We define SC-RDKit score >0.95 as the threshold of SC-RDKit pass rate. **c,** Anchor-distance stress test. Fragment pairs were grouped by anchor distance, and FlexiTAC maintained higher validity across long-distance fragment-pair settings where other 3D models degraded markedly. **d,** Representative long-anchor case generated by FlexiTAC. The model successfully connects a difficult fragment pair with an anchor distance of 26.22 Å. **e,** Fraction of generated PROTACs with linker RMSD below 1.0 and 1.5 Å after force-field optimization. **f,** Hit fraction results under similarity-based recovery thresholds. **g,** Hit-rate curves under different Tanimoto thresholds. **h-i,** Real-world PROTAC rediscovery examples. Given only the fragment inputs, FlexiTAC reconstructs linker designs of Veppanu and PKMYT1 Degrader, demonstrating its ability to recover known PROTAC chemotypes.

Because anchor separation imposes a fundamental geometric constraint on linker design and varies substantially across PROTAC systems, we evaluated 90 fragment pairs spanning anchor distances from 3.1 to 27.1 Å (**Fig. 2c**). This test assesses whether each model can adapt linker length and topology to progressively larger separations between the warhead and E3-ligase ligand. FlexiTAC (ChEMBL) maintained the highest validity across all different anchor distances below 21.1 Å and remained productive in the longest-distance cases. A representative design case at an anchor distance of 26.22 Å is shown in **Fig. 2d**. PocketXMol maintained high validity at shorter anchor distance regions (82.6% in 6.1-9.1 Å, 69.0% in 12.1-15.1 Å), but lost generation capacity when anchor distances in 21.1-27.1 Å, while FlexiTAC (ChEMBL) still maintains 66.1%, 29.6% and 54.4% validity in 21.1-24.0, 24.0-25.3 and 25.3-27.1 Å anchor distance regions, FlexiTAC (scratch) shows even better adaptability in long linker generation, achieving 70.1%, 44.5% and 72.2% validity in 21.1-24.0, 24.0-25.3 and 25.3-27.1 Å anchor distance regions, respectively. This may be due to the distinct distribution of linker lengths, measured by heavy atom count, between the two datasets. The pre-training data contains significantly shorter linkers (range: 3-13, mean: 5.35) compared to the PROTAC-3D dataset (range: 3-40, mean:12.50). Consequently, models trained directly on PROTAC-3D may exhibit better adaptability to longer linkers.

To evaluate the conformational stability of generated PROTACs, we measured the fraction of molecules whose RMSD between the generated and force-field-relaxed conformations stayed below 1.0 Å and 1.5 Å, computed on three test sets (both-unseen, E3-unseen, warhead-unseen). At the stringent 1.0 Å threshold, the two FlexiTAC varia nts consistently ranked first across the three test sets: FlexiTAC(ChEMBL) led on the first set (28.7%), whereas FlexiTAC(scratch) performed best on the second and third s ets (26.3% and 23.6%, respectively). The best-performing baseline, PocketXMol, reac hed 21.8%, 15.0% and 10.6%, less than half of the fraction on the warhead-unseen set. DiffLinker and LinkerNet remained below 18.1% and 10.4% on every set, while most PROTACs provided by 3DLinker failed because it lacked the ability to generate high-quality linker conformations for force-field relaxation. At the 1.5 Å threshold, FlexiTAC(ChEMBL) outperformed PocketXMol by 2.0, 17.6 and 22.1 percentage points on the both-unseen, E3-unseen and warhead-unseen sets, respectively. FlexiTAC(scratch) similarly exceeded PocketXMol by 17.2 and 26.2 percentage points on the E3-unseen and warhead-unseen sets, while trailing it by only 0.9 percentage points on the both-unseen set. (**Fig. 2e**).

Together, these results indicate that ChEMBL3D pretraining provides transferable chemical and geometric priors, modestly improving molecular validity, atom stability and robustness across unseen fragment pairs and common anchor separations. This advantage diminished at the longest separations, where FlexiTAC(scratch) performed bet ter, consistent with the substantially shorter linkers represented in the pretraining corpus. Thus, pretraining enhances general fragment-connection quality, whereas robust long-range connection remains dependent on sufficient coverage of PROTAC-relevant linker lengths. Together, these results demonstrate that FlexiTAC generates chemically valid PROTACs with geometrically plausible conformations that remain stable after force-field relaxation across diverse fragment conditions.

### FlexiTAC achieves robust PROTAC rediscovery and chemical-space coverage across unseen fragment conditions

Beyond basic molecular-quality metrics, the ability to rediscover known bioactive PROTACs provides a more direct measure of practical design potential. We therefore assessed whether FlexiTAC could recover reference PROTACs from their constituent fragment pairs using two complementary metrics. Hit rate measures the proportion of generated molecules that match a reference PROTAC, whereas hit fraction measures the proportion of reference PROTACs recovered by at least one generated molecule. Strict recovery was evaluated at a molecular-fingerprint similarity of 1.0, and similarity-based recovery was assessed at progressively lower similarity thresholds.

Both FlexiTAC variants provided strong coverage of the reference chemical space (**Fig. 2f**). In the E3-unseen test set, FlexiTAC(ChEMBL) and FlexiTAC(scratch) achieved strict hit fractions of 7.60% and 7.13%, respectively, compared with 3.62% for the strongest non-FlexiTAC baseline, Link-INVENT. At Tanimoto >0.9, their hit fractions increased to 19.25% and 22.61%, exceeding the second-best baseline by 7.36 and 10.72 percentage points, respectively. In the warhead-unseen test set, the two variants achieved strict hit fractions of 2.54% and 3.71%, compared with 1.36% for Link-INVENT. At Tanimoto >0.9, FlexiTAC(ChEMBL) and FlexiTAC(scratch) reached 14.98% and 22.08%, outperforming the strongest baseline, DiffLinker by 6.34 and 13.44 percentage points, respectively. The advantage was also maintained in the both-unseen setting: the two variants recovered 5.73% and 7.82% of the references under the strict criteria, compared with 3.88% for Link-INVENT. At Tanimoto >0.9, FlexiTAC(scratch) achieved the highest hit fraction of 24.91%, exceeding Link-INVENT by 7.64 percentage points.

The hit rate results further showed that FlexiTAC efficiently generated reference-l ike PROTACs (**Fig. 2g**). At Tanimoto = 1.0, in the both-unseen test set, FlexiTAC(ChEMBL) and FlexiTAC(scratch) achieved hit rates of 10.63% and 6.30%, respectively, compared with 3.03% for the strongest baseline, DiffLinker. At Tanimoto > 0.9, the tw o variants reached nearly identical hit rates of 25.17% and 25.20%, exceeding the strongest baseline, Link-INVENT by more than 16 percentage points. In the E3-unseen test set, their strict hit rates were 6.06% and 5.90%, compared with 2.94% for LinkerNet. At Tanimoto similarity>0.9, FlexiTAC(ChEMBL) reached 9.94% and FlexiTAC(scr atch) reached 13.56%. These rates exceeded LinkerNet’s 4.24% by 5.70 and 9.32 percentage points, respectively. In the warhead-unseen setting, FlexiTAC(ChEMBL) and FlexiTAC(scratch) achieved hit rates of 4.08% and 6.12%, followed by 15.80% and 2 0.50% at Tanimoto >0.9. The latter results exceeded LinkerNet by 5.32 and 10.02 percentage points.

At the more permissive Tanimoto >0.8 threshold, which reflects recovery across a broader region of related chemical space, both FlexiTAC variants remained ahead of the baselines in all three test sets. FlexiTAC(ChEMBL) and FlexiTAC(scratch) achieved hit rates of 39.04% and 44.40% in E3-unseen, compared with 26.78% for Link-INVENT. On warhead-unseen, FlexiTAC(ChEMBL) reached 41.28% and FlexiTAC(scratch) reached 50.64%, compared with 34.24% for LinkerNet. On both-unseen, FlexiTAC(ChEMBL) achieved 51.47% and FlexiTAC(scratch) achieved 48.63%, versus 34.30% for Link-INVENT. These results demonstrate its ability to recover known linkers while efficiently exploring closely related chemical space. We further tested FlexiTAC with two realistic rediscovery cases. As demonstrated in **Fig. 2h,i**, the first used the FDA approved PROTAC degrader Veppanu. The second used the fragment information from a PKMYT1 Degrader^14^. In both cases, FlexiTAC was given the two fragments as condition, and successfully generated linkers that are exactly identical to those in the original PROTACs.

Additionally, we provide evaluation results for FlexiTAC trained on the PROTAC-DB 1.0 and ZINC datasets to compare it with prior baselines(**Supplementary Table-1** and **Supplementary Table-2**.). This comparison further demonstrates that FlexiTAC’s superiority is not solely attributable to the scale of the training data. Across both datasets, FlexiTAC consistently outperforms previous models in validity and conformational stability. Specifically, on the ZINC dataset, FlexiTAC achieves the highest validity (98.9 ± 0.0%) and recovery rate (33.9 ± 0.9%), while delivering the lowest RMSD (1.36 ± 0.0 Å) and minimum energy (E_min_ of 23.6 ± 0.1 kcal/mol) after force-field relaxation. Similarly, on PROTAC-DB 1.0, it maintains superior conformational quality with the lowest RMSD (1.37 ± 0.04Å) and significantly improved recovery (23.3 ± 0.0%).

Together, the benchmark and case-study results establish FlexiTAC as a high-fidelity linker generator capable of recovering established active PROTACs and related chemotypes under challenging unseen-fragment conditions. This capacity to prioritize application-relevant linker hypotheses could accelerate PROTAC discovery.

### FlexiTAC learns fragment-specific linker preferences across unseen conditions

For practical PROTAC discovery, a linker-generation model must respond to the chemical and geometric information encoded by its input fragments^31^. However, a strong general linker prior could recover known chemotypes across diverse inputs without preferentially generating them for their cognate fragment pairs. Such condition-insensitive recovery would limit the model’s utility for rational, fragment-guided PROTAC design. Although FlexiTAC achieved high hit rates and broad reference recovery in the preceding analyses, these results alone do not establish fragment awareness. We therefore introduced the Fragment-Aware Score (FAScore) to measure how strongly linker recovery depends on the supplied fragment pair. For each fragment pair, FAScore compares the linker recovery rate under the current fragment pair condition with the background recovery rate across all fragment conditions. A FAScore of at least 10 indicates that a reference linker is recovered at least tenfold more frequently under its corresponding fragment condition than in the global background.

FlexiTAC showed the broadest coverage of strongly enriched fragment pairs across all three test sets (**Fig. 3**). In the both-unseen setting, 66.7% of fragment pairs achieved FAScore ≥10 with FlexiTAC (scratch), compared with 60.0% for FlexiTAC (ChEMBL) and 26.7% for DiffLinker, the strongest baseline in this test set. For the top-3 model, the corresponding proportions were 46.0% of FlexiTAC(scratch), 42.0% of FlexiTAC(ChEMBL) and 26.0% of Link-INVENT in the E3-unseen set. In the warhead-unseen set, these proportions were 32.0% of FlexiTAC(scratch), 30.0% of FlexiTAC(ChEMBL) and 20.0% of LinkerNet, respectively.

**Figure. 3.**
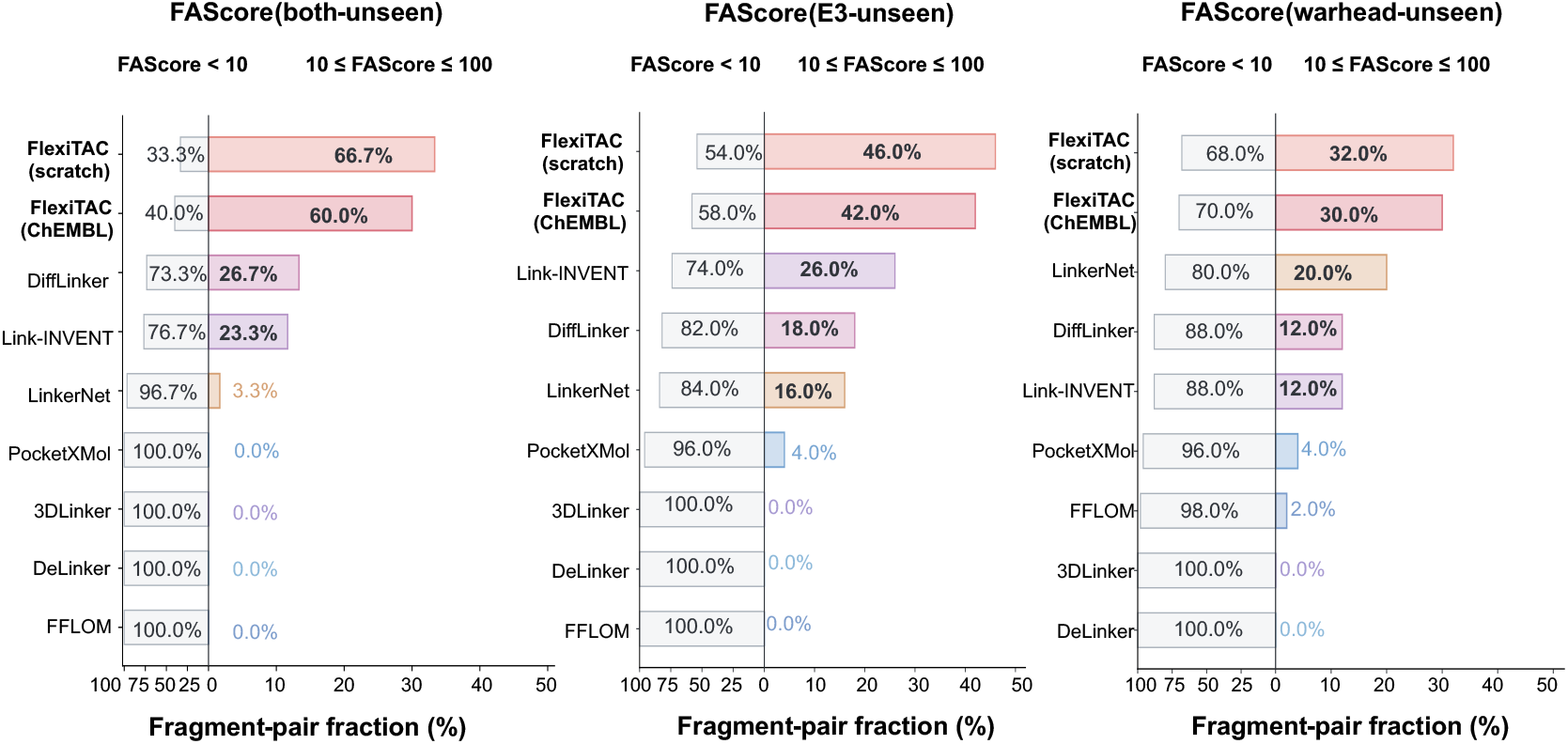
| Fragment-aware generation performance of PROTAC linker design models. Fragment-pair fractions are shown for the both-unseen, E3-unseen and warhead-unseen test sets. For each model, fragment pairs were divided into those with low fragment awareness (FAScore < 10; left) and those showing strong fragment-awareness (10 ≤ FAScore ≤ 100; right). Higher scores therefore indicate that linker recovery is more strongly associated with the supplied fragment condition. Percentages denote the fraction of fragment pairs within each FAScore interval.

Across all 130 fragment pairs, FlexiTAC (scratch) and FlexiTAC (ChEMBL) achieved 10 ≤ FAScore ≤ 100 for 59 and 54 pairs, respectively. By comparison, Link-INVENT, the strongest baseline overall, reached this interval for only 26 pairs. DiffLinker showed the strongest overall fragment-aware performance among the evaluated 3D baselines, but its coverage remained substantially below that of either FlexiTAC variant. FlexiTAC(scratch), trained exclusively on PROTAC-3D, exhibited slightly stronger fragment-conditioned enrichment than the ChEMBL-pretrained variant. This difference may reflect the broader chemical prior acquired during pretraining, which could modestly weaken specialization to PROTAC-specific fragment–linker relationships. Nevertheless, both variants substantially outperformed all baselines, confirming that FlexiTAC generates linkers in a manner responsive to the input fragment conditions. Link-INVENT performed comparatively well among the 2D baselines but remained substantially below FlexiTAC, suggesting that two-dimensional representations of chemical identity and molecular connectivity alone may be insufficient to capture fragment-conditioned enrichment. This performance gap supports FlexiTAC’s central design principle of jointly modelling linker chemistry and three-dimensional geometry to learn fragment–linker compatibility.

### FlexiTAC accurately recovers local geometry and generates physically plausible PROTAC conformations

Beyond generating chemically valid PROTACs, an effective linker model must accurately reproduce the geometric patterns required for compatibility with the terminal fragments. To evaluate this, we compared the linker geometry distributions generated by each model against those of reference molecules. This comparison was quantified using the Jensen-Shannon divergence (JSD) across seventeen local motif types, encompassing nine bond-length types, six bond-angle types, and two torsional motifs. **Fig. 4a** presents Gaussian-smoothed density profiles for the nine representative torsion-angle, bond-angle and bond-length motifs, suitable for observing peak positions, multimodal features, and overall distribution shifts. This advantage was particularly pronounced for chemically informative motifs, such as C-N bonds (JSD of 0.444 vs 0.461-0.831 for baselines) and C-N-C angles (0.402 vs 0.478-0.806). Across the evaluated dihedral motifs, both FlexiTAC variants more closely reproduced the motif-specific reference distributions than the baseline models. They retained multiple modes when these occurred in the reference while also recovering the predominantly unimodal distribution of other motifs. Several baselines instead broadened distinct modes or merged them into a single peak. These results indicate that FlexiTAC captures motif-dependent dihedral preferences rather than imposing a common distribution shape. Additionally, FlexiTAC(ChEMBL)’s mean JSDs for bond angles, bond lengths and torsion angles were 0.471, 0.556 and 0.557, respectively, compared with 0.498, 0.569 and 0.596 for FlexiTAC(scratch), indicating that pretraining on quasi-PROTAC data provided an additional gain in conformational fidelity, enabling more accurate recovery of local geometric preferences across diverse chemical environments. **Supplementary Fig. 6** reports unsmoothed distributions for a broader set of geometric motifs.

**Figure 4.**
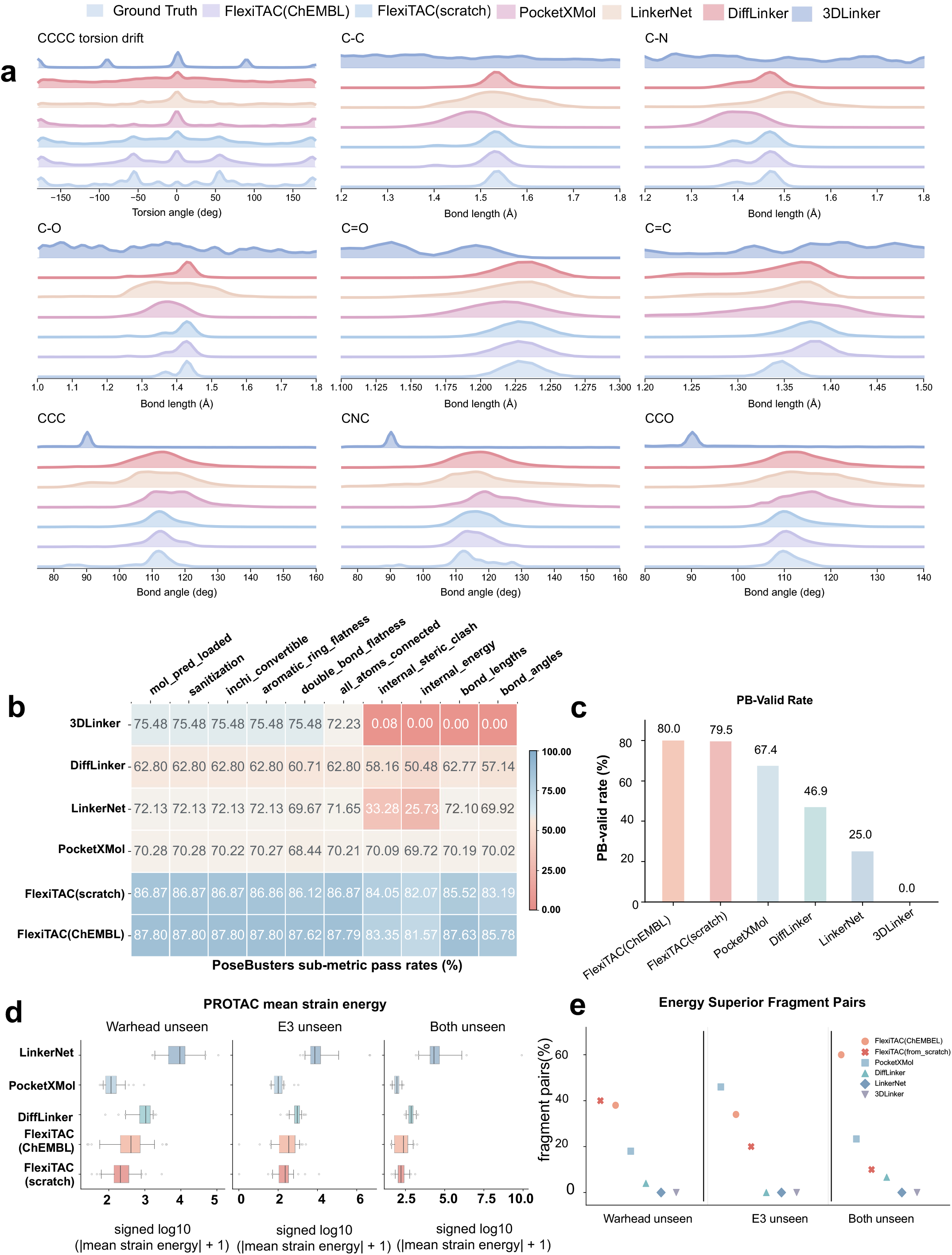
| Geometric fidelity and conformational quality of generated PROTAC linkers. **a,** Distribution drift figures for nine linker geometry substructures, including torsion angles, bond lengths, and bond angles. A distribution that is more similar to the reference drift figures of reference PROTACs indicates a better distribution learned by the model. **b,** Detailed PoseBusters sub-metric pass rates for generated molecules. **c,** Overall PoseBusters pass-all rate for the linker structures generated by models. **d,** Strain energy distributions across the warhead-unseen, E3-unseen, and both-unseen test sets. Lower strain energy indicates better PROTAC conformation. **e,** Fraction of fragment pairs for which each model had the smallest mean energy decrease after MMFF94 minimization.

We then applied PoseBusters checks to assess whether this distributional fidelity translated into physically reasonable 3D linker conformations. As shown in **Fig. 4b,c**, both FlexiTAC variants achieved the highest overall Posebusters pass-all(PB-valid) rates among the compared models (80.0% for the FlexiTAC(ChEMBL) and 79.5% for FlexiTAC(scratch)), surpassing PocketXMol by 12.6 and 12.1 percentage points, respectively, whereas several baselines failed specific geometric checks (for instance, 3DLinker passed virtually none of the checks, yielding a PB-valid rate of 0.0%) or showed markedly lower final pass rates. Collectively, FlexiTAC not only modeled local geometric fidelity with high accuracy but also achieved substantial gains in overall geometric modeling.

We assessed the energetic quality of complete PROTAC structures by measuring PROTAC strain energy across the three test sets. FlexiTAC maintained low strain energy distributions in the three held-out settings. Across the three held-out settings, FlexiTAC (scratch) achieved median PROTAC strain energies of 155.80, 227.20, and 207.55 kcal/mol, respectively, and the FlexiTAC (ChEMBL) yielded medians of 172.72, 345.66, and 410.72 kcal/mol on the corresponding test set. Both variants remained markedly below DiffLinker (692.92, 943.37 and 1006.54 kcal/mol) and LinkerNet (20655.98, 7380.81, and 9589.69 kcal/mol); PocketXMol attained the lowest medians (89.33, 100.34, and 116.57 kcal/mol), potentially reflecting the energetic prior acquired from its extensive small-molecule pretraining.

At the fragment-pair level, we calculated the energy difference of the model-gener ated PROTACs before and after force-field relaxation to quantify the energy stability. For a given fragment-pair, the model that achieved the lowest mean energy change wa s considered to outperform the others. As shown in **Fig. 4e**, the FlexiTAC(ChEMBL) won the largest overall share of fragment pairs across the benchmark (54 of 130 pairs), including 60% on the both-unseen test set, 34% on the E3-unseen test set, and 38% on the warhead-unseen test set. FlexiTAC(scratch) achieved 10%, 20%, and 40% on the corresponding splits, leading the warhead-unseen set, while PocketXMol led narrowly on the E3-unseen set (46%). The series of conformation-related metrics demonstrates that FlexiTAC better preserves linker-level geometric distributions and produces more physically acceptable 3D conformations than current 3D baselines.

These complementary results clarify the effects of pretraining. Training exclusively on PROTAC-3D produced lower absolute strain energies, indicating that the curated PROTAC-specific data already provide a strong energetic prior and may better represent the size and linker-length distributions of full PROTACs. ChEMBL3D pretraining did not further reduce absolute strain energy but substantially increased the number of fragment pairs requiring minimal energetic adjustment during relaxation, particularly i n the both-unseen setting, indicating improved transferability of local geometric priors. Thus, pretraining enhances geometric robustness and relaxation stability, although its energetic benefit is constrained by the distributional mismatch between quasi-PROTAC pretraining molecules and PROTAC-specific chemical space.

### FlexiTAC remains robust as fragment-pair geometric difficulty increases

The performance of structure-based linker design models is affected not only by whether the fragment-pair condition was present in the training set, but also by the varying levels of difficulty arising from the geometric conformation of the fragment pairs. Therefore, we first analyzed the three test sets of PROTAC-Bench using four structural variables: anchor distance, total fragment size, linker ring count and linker atom count (**Fig. 5a-d**). The warhead-unseen set presented the strongest geometric constraints, with a mean anchor distance of 17.1 Å and 21.1% of reference PROTACs exceeding 20 Å, compared with 14.1 Å and 2.5% for E3-unseen, and 11.3 Å and 11.7% for both-unseen. It also contained the longest linkers, with a mean of 15.8 atoms and 16.1% containing more than 20 linker atoms. By comparison, these proportions were 10.5% for E3-unseen and 4.5% for both-unseen. E3-unseen was instead enriched in structurally complex linkers: 76.2% contained more than one ring and 29.0% contained more than two rings, compared with 60.9% and 15.0% for both-unseen,75.1% and 26.5% for warhead-unseen, respectively. The mean total fragment sizes were similar across the three sets, ranging from 49.6 to 52.6 atoms, indicating that their differences arose primarily from linker topology and fragment separation rather than a simple shift in fragment size.

**Figure 5.**
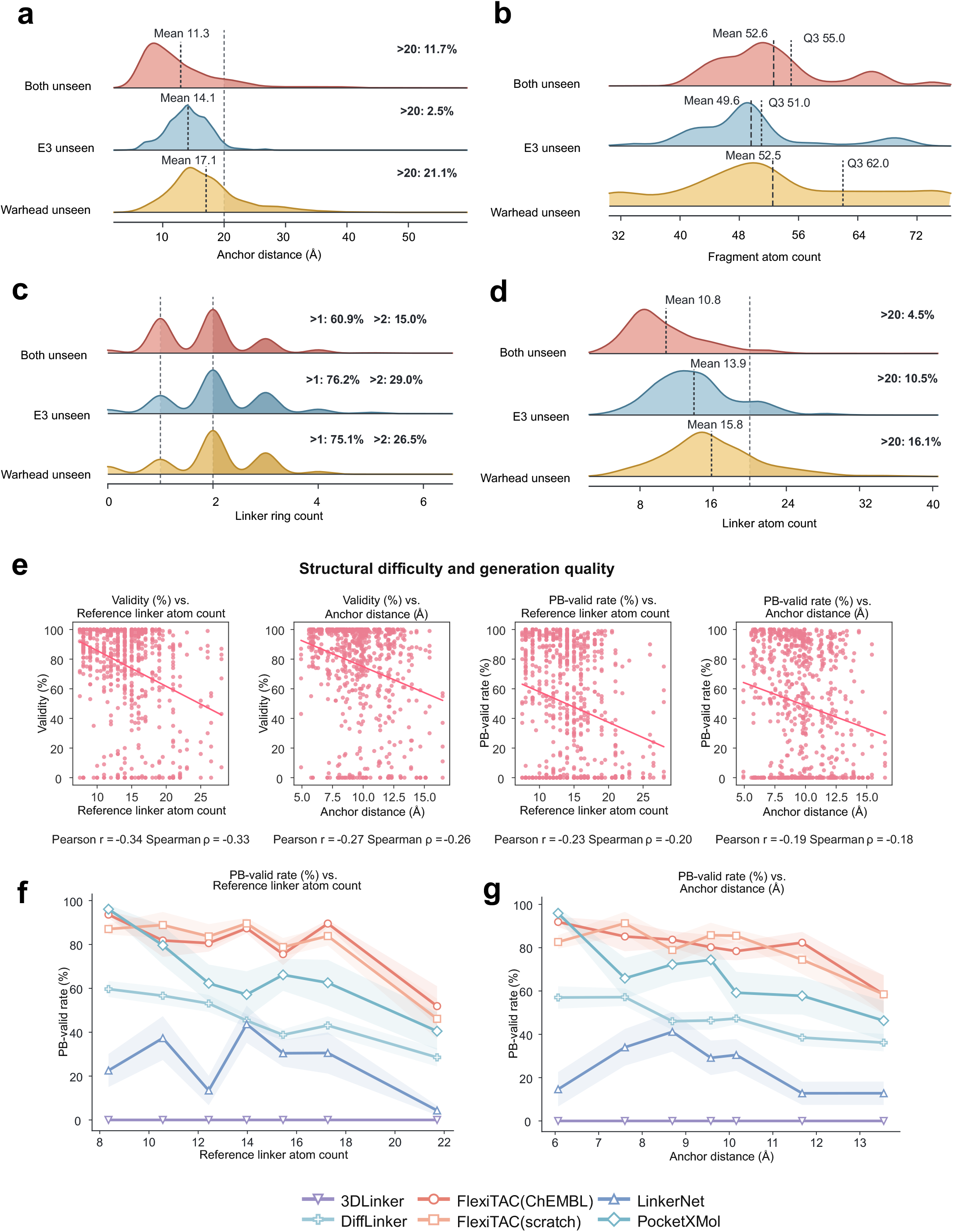
| Correlation analysis of geometric constraints and model performance. **a**, Distribution of anchor distances between the two attachment atoms across the three test sets; dashed lines indicate the mean and 20 Å thresholds. **b**, Distribution of the combined atom count of the two input fragments; dashed lines indicate the mean and third quartile. **c**, Distribution of linker ring counts, with annotations showing the proportions containing more than one or more than two rings. **d**, Distribution of linker atom counts; dashed lines indicate the 20-atom threshold, and annotations report the proportion above this threshold. **e**, Relationships between reference linker atom count or anchor distance and validity or PB-valid rate across all 3D linker-design models and 130 fragment-pair systems. Solid lines show linear fits; Pearson and Spearman correlation coefficients are reported below each panel. **f**, Correlation of PB-valid rate and reference linker atom count. **g**, Correlation of PB-valid rate and anchor distance.

We next examined whether these structural constraints were associated with generation performance across the six evaluated 3D linker-design models (**Fig. 5e**). The pooled analysis comprised 130 fragment-pair systems for each model, giving 780 model-system observations. Reference linker length showed the strongest association with validity, with Pearson (r= −0.34) and Spearman (*ρ* = −0.33). Anchor distance was also negatively associated with validity (r= −0.27, *ρ* = −0.26). The corresponding relationships with PB-valid rate were weaker but remained consistently negative: *r* = −0.23 and *ρ* = −0.20 for linker length, and *r* = −0.19, *ρ* = −0.18 for anchor distance. These results indicate that more complex linkers and more widely separated anchors reduce both the probability of generating a connected, chemically valid PROTAC and the probability that its reconstructed linker satisfies all PoseBusters checks.

To resolve how individual models responded to increasing geometric difficulty, we divided the 130 fragment pairs into seven ranges according to the number of reference linker atoms or anchor distance and calculated the mean PB-valid rate within each range (**Fig. 5f,g**). PB-valid rates generally declined as either constraint increased. Across linker atom count bins, FlexiTAC(ChEMBL) decreased from 93.6% for the shortest-linker range, with 8.4 atoms on average, to 51.9% for the longest-linker range, with a mean of 21.7 atoms. FlexiTAC(scratch) decreased from 87.0% to 46.1% over the same range. In the longest-linker bin, both variants of FlexiTAC remained ahead of PocketXMol (40.5%), DiffLinker (28.5%), LinkerNet (4.3%) and 3DLinker (0%). A similar pattern was observed for anchor separation. In the most widely separated bin, with a mean anchor distance of 13.5 Å, FlexiTAC(ChEMBL) and FlexiTAC(scratch) retained PB-valid rates of 58.4% and 58.5%, respectively, compared with 46.3% for PocketXMol, 36.2% for DiffLinker and 12.8% for LinkerNet. Consistent with these trends, FlexiTAC retained a clearer advantage as geometric constraints increased. These results reveal a clear dependence of linker-generation performance on fragment-pair difficulty and demonstrate the robustness of both FlexiTAC variants across diverse geometric conditions.

### FlexiTAC efficiently explores PROTAC-relevant chemical space at inference time

In practical screening, we assume that a PROTAC linker design model should preserve the ability to explore a large conditional design space, preserve the geometry imposed by two fragments and return high-quality candidates within a limited sampling budget. FlexiTAC addresses this requirement by using BFN to model continuous 3D coordinates and discrete atom types in a shared continuous parameter space. This method improves sampling speed and enables the model to better capture the data manifold and reduce error, which is important for linker design because small errors in atom identity or geometry can affect linker’s overall chemical identity.

We first examined the sampling trajectory of FlexiTAC for a representative fragment pair, tracking the evolution of linker atom identities and 3D geometry to assess convergence in chemical composition and molecular structure (**Fig. 6a**). Given that FlexiTAC (ChEMBL) exhibits high effectiveness, PB-valid rate, and superior performance in geometry substructures, it was adopted as the default choice for subsequent experiments. In the 100-step sampling trajectory, atom types changed primarily during the early part of generation, of which 95.9% occurred within the first 50 steps. After this point, the atom-type became nearly stable, indicating that the later updates mainly refine geometry rather than repeatedly changing chemical composition. We then compared structural convergence with other 100-step generative models using the linker RMSD to the final generated conformation (**Fig. 6b**). FlexiTAC showed the fastest decline in mean linker RMSD, decreasing from 2.91 Å at the beginning of sampling to 0.49 Å at step 50. In contrast, DiffLinker and PocketXMol retained higher intermediate RMSD at step 50, 1.89 Å and 1.44 Å, respectively. The case in **Fig. 6d** shows the same behavior visually, with isolated atoms at the beginning, partial linker organization by step 25 and nearly stable halfway through inference. This faster convergence is consistent with the design and expectation of the BFN’s parameter space setting.

**Figure 6.**
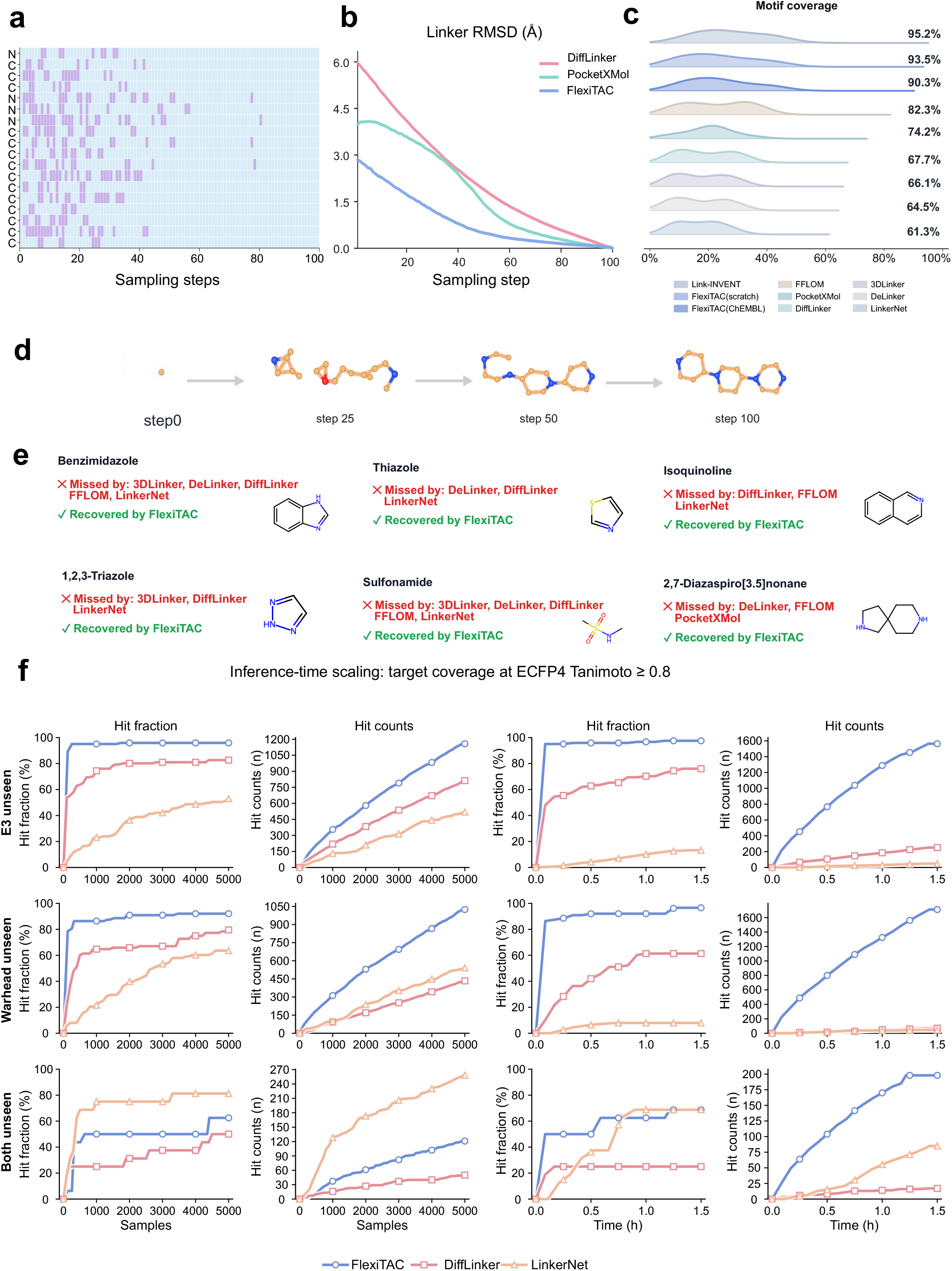
| Sampling dynamics, structural coverage and design efficiency of FlexiTAC. **a,** FlexiTAC’s sampling trajectory for a representative linker. Blue indicates that the predicted atom type remained unchanged from the preceding step, whereas purple marks a change in the predicted atom type. Atom types change frequently during the early sampling steps and soon become stable after half of the sampling process; this behavior is consistent with the Bayesian-flow formulation, in which uncertainty is progressively reduced during sampling. **b,** Linker RMSD trajectories compare FlexiTAC with other 100-step 3D generative models. FlexiTAC converges faster, indicating efficient use of sampling steps for structural refinement. **c,** Motif coverage rate of generated linkers. We extracted 62 distinct linker substructures from the reference PROTAC test sets and evaluated whether standard sampled molecules from each model on the benchmark successfully covered these motifs. **d,** A representative case of FlexiTAC’s 100-step linker generation process. **e,** Coverage of specific common linker substructures from the test set. FlexiTAC successfully recovers several frequent motifs that are missed by baseline models, suggesting that it effectively learns chemical patterns instead of generating generic chains. **f,** Inference-time scaling analysis of target coverage across different test sets. We treated generated PROTACs with ECFP4 fingerprint similarity ≥ 0.8 as the hit targets. The metrics evaluated include the hit fraction and hit counts (the number of unique generated molecules that matched at least one reference after deduplication). The left two columns illustrate the scaling performance as the total number of sampled molecules increases, while the right two columns demonstrate the performance as a function of extended inference time.

Structural-motif recovery measures whether generated linkers capture recurring chemical features of known PROTACs. We therefore evaluated motif-coverage rates using the 62 references motifs found in the PROTAC-Bench test-set linkers, including common functional groups and recurrent ring systems (**Supplementary Fig. 7 and 8**). Details regarding the curation of reference motifs are provided in the Methods section. A motif was considered recovered when it matched at least one valid generated linker by RDKit substructure matching. FlexiTAC showed the strongest coverage among the evaluated 3D models. As shown in **Fig. 6c**, FlexiTAC(scratch) recovered 58 of 62 motifs (93.5%), ranking second across all models after Link-INVENT (59 of 62; 95.2%) and performed best among all structure-based baselines. FlexiTAC (ChEMBL) recovered 56 motifs (90.3%). By comparison, PocketXMol, DiffLinker, 3DLinker and LinkerNet achieved coverage rates of 74.2%, 67.7%, 66.1% and 61.3%, respectively.

Representative cases of motif coverage are shown in **Fig. 6e**. FlexiTAC recovered benzimidazole and sulfonamide motifs that were missed by five baseline methods. It also recovered thiazole and 1,2,3-triazole motifs that were absent from several baseline outputs. Both FlexiTAC variants further captured the 2,7-diazaspiro[3.5]nonane scaffold, which was missed by DeLinker, FFLOM and PocketXMol. These results indicate that FlexiTAC learned a broad repertoire of linker substructures, spanning aromatic heterocycles, polar functional groups and conformationally constrained spirocyclic motifs.

To test whether FlexiTAC’s generative performance translated into efficiency in PROTAC design, we constructed three inference-time scaling tasks from PROTAC-Bench (**Fig. 6f**). We selected one fragment pair from each test set that contained the greatest number of PROTACs, and treated all associated PROTACs with the same fragment pair as reference targets. The resulting sets contained 121 E3-unseen, 88 warhead-unseen and 16 both-unseen PROTACs. Generated molecules were compared with these targets using full-molecule ECFP4 fingerprints. A reference was considered recovered when its fingerprint similarity to at least one generated molecule was ≥ 0.8, which measures the recovery of nearby chemical space. Hit fraction denotes the proportion of unique reference PROTACs recovered. After removing duplicate generated molecules within each sampling budget, generated hit count denotes the number of unique generated molecules matching at least one reference. We selected FlexiTAC(ChEMBL) for this comparison because it achieved higher validity and PoseBusters pass rates, making it suitable for large-scale generation experiments.

Within the 5,000-sample budget, FlexiTAC recovered 116 E3-unseen references (95.9%) and 81 warhead-unseen references (92.0%). It produced 1,158 and 1,024 unique generated hits in these tasks, respectively. These values exceeded the strongest baseline by 345 hits for E3-unseen and 483 hits for warhead-unseen. The both-unseen task was more challenging. FlexiTAC recovered 10 of 16 references (62.5%) and produced 121 unique generated hits. This exceeded the 50 hits produced by DiffLinker but remained below the 258 hits produced by LinkerNet.

Although LinkerNet achieved a higher generated hit count for this pair, this pair-specific recovery did not translate into broad chemical exploration. LinkerNet showed only 21.1% uniqueness and recovered 61.3% of the test-set linker motifs. By comparison, FlexiTAC(ChEMBL) and FlexiTAC(scratch) achieved motif coverages of 90.3% and 93.5%, respectively. They also maintained higher uniqueness values of 55.1% and 51.8%. Thus, although FlexiTAC was not the strongest method for every individual fragment pair, it sampled a broader and less redundant region of linker chemical space. This broader exploration may increase the likelihood of identifying novel and structurally diverse candidates rather than repeatedly generating a narrow set of chemotypes. For the time-based curves, FlexiTAC had already achieved a hit fraction of 95.0% on E3-unseen, 86.4% on warhead-unseen and 50% on both-unseen test set after only 0.085 h. These early samples contained 214, 223 and 25 PROTACs that recover references, respectively. Recovery continued to increase with runtime. Within 1.5 h, FlexiTAC reached hit fractions of 97.5%, 96.6% and 68.8% across the three tasks, and produced 1,563 E3-unseen, 1,711 warhead-unseen and 198 both-unseen hits, which largely surpassd the baseline models.

Within the same time budget, these generated hit counts were 6.2-fold, 24.4-fold and 2.3-fold higher than those of the strongest competing method in the respective tasks. FlexiTAC also recovered 26 more E3-unseen references and 31 more warhead-unseen references than DiffLinker. In the both-unseen task, FlexiTAC and LinkerNet each recovered 11 of 16 references. However, FlexiTAC generated 198 matching molecules compared with 85 from LinkerNet. Thus, although both-unseen recovery remained more difficult, FlexiTAC’s higher sampling throughput yielded a substantially larger pool of chemically relevant candidates within a fixed time budget.

Together, the combination of rapid convergence, broad motif coverage and efficient reference recovery supports FlexiTAC as a practical method for PROTAC linker design. Effective linker design requires not only rapid sampling, but also sufficient exploration of chemically relevant and structurally compatible linker space; FlexiTAC balances these requirements by recovering diverse PROTAC-like chemotypes within limited sampling budgets.

### Programmable control of linker conformational flexibility via CERScore and posterior guidance

Linker flexibility is not well captured by a single topological descriptor of a molecule. We therefore defined a conformational ensemble-derived rigidity score (CERScore) *r*_rigid_ (**Fig. 7a**). For each linker, we considered three complementary sources of motion: we generated 100 conformations for each PROTAC, and calculated the variance in the distance between the two fragment anchor atoms, the number of rotatable bonds in the linker, and the variation in intra-linker distances for atom pairs separated by more than two bonds along the linker topology. These terms respectively measure how freely the two fragments can move relative to each other, how many internal rotations the linker allows, and whether the long-range linker geometry is preserved across conformations. The final score is a weighted combination of these components where a larger value indicates a more rigid linker. The details of rigidity score construction is described in Methods.

**Figure 7.**
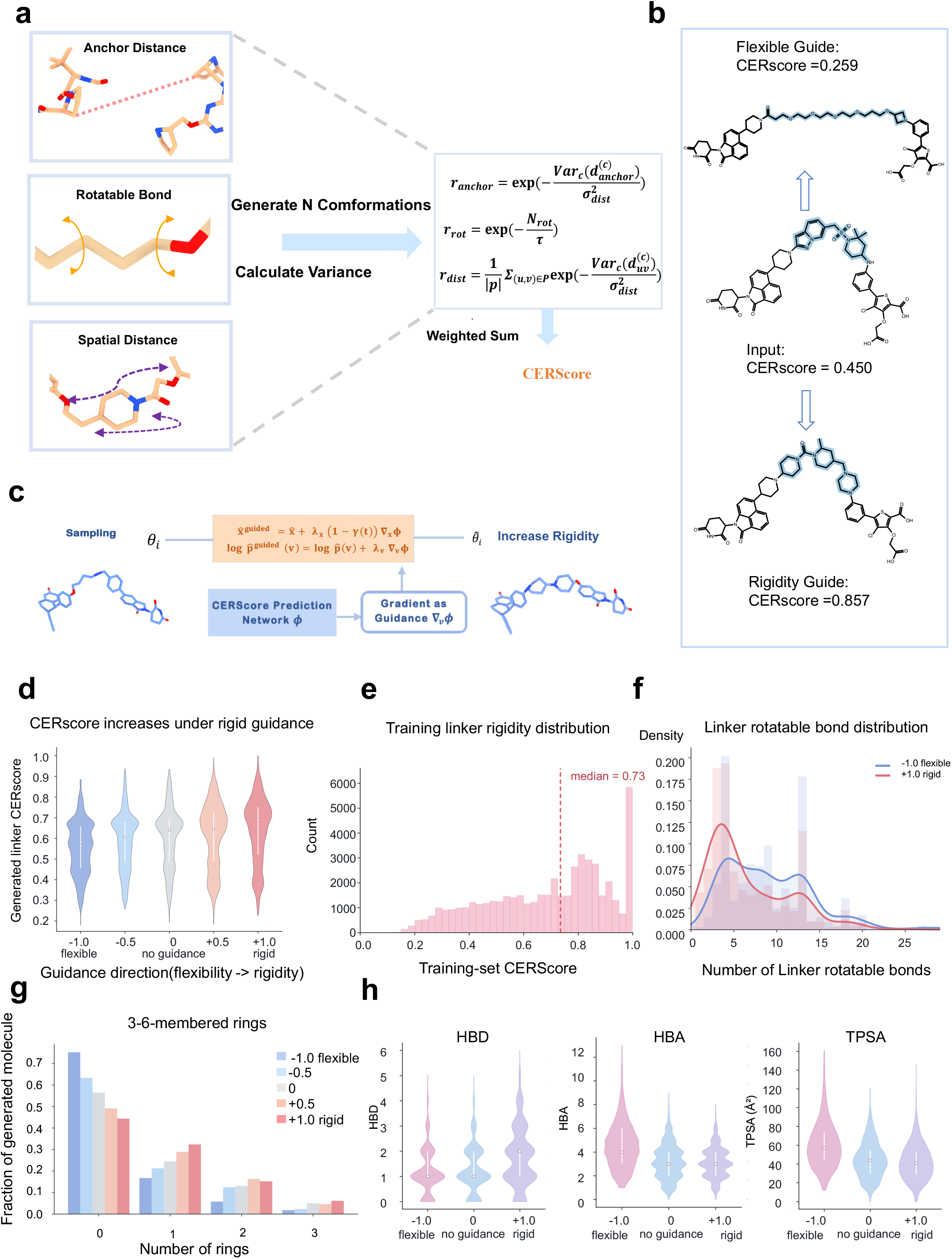
| Controllable linker rigidity through posterior gradient guidance. **a,** Construction of linker CERScore from PROTAC structures. **b,** Representative molecules generated from the same fragment input under flexible, unguided and rigid settings, showing that the guidance changes linker shape and rigidity within the same fragment context. **c,** The posterior gradient-guidance module can steer sampling toward a desired linker rigidity while receiving the same fragment-pair context. **d,** Distribution of generated CERscores under different guidance strengths. The distribution shifts monotonically from flexible to rigid guidance, confirming controllable sampling. **e,** Training-set rigidity distribution used to define the target range for guidance. **f, g, h,** Structural-property analysis under guidance. Rigid guidance reduces rotatable-bond distributions and changes the 3-6-membered ring distribution, while HBD, HBA and TPSA remain within a chemically reasonable range. These results show that FlexiTAC can tune linker flexibility without collapsing molecular quality.

A lightweight predictor maps the final backbone embeddings to this CERscore. During the final 60 sampling steps, the gradient of the predictor biases coordinates and atom-type parameters. The generator remains fixed, so guidance changes the sampling trajectory rather than retraining the backbone. This design keeps the generation process to the learned fragment-conditioned distribution while allowing a user-specified direction and intensity of control.

To evaluate controllability, we first assigned rigidity scores to the PROTAC-Bench test sets and then selected ten test PROTACs spanning the low-to-high rigidity range. For each corresponding fragment pair, we sampled 100 linkers under five guidance strengths, from flexible guidance (-1.0) to rigid guidance (+1.0). The molecular examples in **Fig. 7b** illustrate the intended effect: starting from an input design with a rigidity score of 0.450, flexible guidance produces a more extended and flexible linker with a lower score (0.259), whereas rigid guidance produces a more compact, ring-rich linker with a higher score (0.857). More cases are provided in **Supplementary Figs. 9,10**. Across these fragment pairs, the mean predicted score increased from 0.554 to 0.635 as the guidance weight changed from -1.0 to +1.0. The median increased from 0.565 to 0.670 (**Fig. 7d**).

The guidance-induced shifts in rigidity scores were accompanied by chemically interpretable structural changes. To interpret the structural basis of flexibility control, we first examined the distribution of rigidity scores in the training set, which had a mean of 0.73 ((**Fig. 7e**). Flexible guidance enriched linkers with more rotatable bonds, whereas rigid guidance shifted the distribution towards fewer rotatable bonds, with the mean count decreasing from 9.12 at a guidance strength of -1.0 to 6.67 at +1.0 (**Fig. 7f**). In parallel, the distribution of 3-6 membered rings moved in the opposite direction (**Fig. 7g**). The fraction of generated linkers containing no rings decreased from 75.2% to 44.4%, while the fraction containing one such ring increased from 16.8% to 32.4%, the fraction containing two rings increased from 5.8% to 15.2% and the fraction containing three rings increased from 1.8% to 6.2%. The guidance therefore changed interpretable structural features, not only the predictor output.

Rigidity guidance also induced interpretable shifts in linker physicochemical properties (**Fig. 7h**). Flexible guidance favoured heteroatom-rich chains, increasing median HBA count and TPSA to 4 and 56.8 Å², whereas rigid guidance reduced these values to 3 and 41.1 Å² and modestly increased the median HBD count from 1 to 2, consistent with enrichment of constrained ring-containing scaffolds. The distributions remained broad, indicating that guidance modulated linker polarity and hydrogen-bonding capacity without collapsing chemical diversity.

### FlexiTAC Enables Structure-based PROTAC and Macrocyclic Linker Design from Experimental and Computationally Predicted Complexes

SBDD for PROTACs requires linker generation from structural inputs that vary substantially in source, completeness and uncertainty. Benchmark studies usually provide predefined fragment poses, whereas practical discovery campaigns often lack experimentally resolved ternary complexes. When a ternary structure is available, the warhead and E3-ligase ligand poses can be extracted directly. Otherwise, the fragments must be redocked into experimental protein structures or into computationally predicted POI-E3 complexes. We therefore evaluated FlexiTAC across these three SBDD scenarios (**Fig. 8**). We further examined a macrocycle-like PROTAC to assess generation beyond conventional linear, two-anchor linker topologies. Together, these experiments tested whether FlexiTAC could extend fragment-conditioned linker generation from standardized benchmarks to practical SBDD settings with varying levels of structural information.

**Figure 8.**
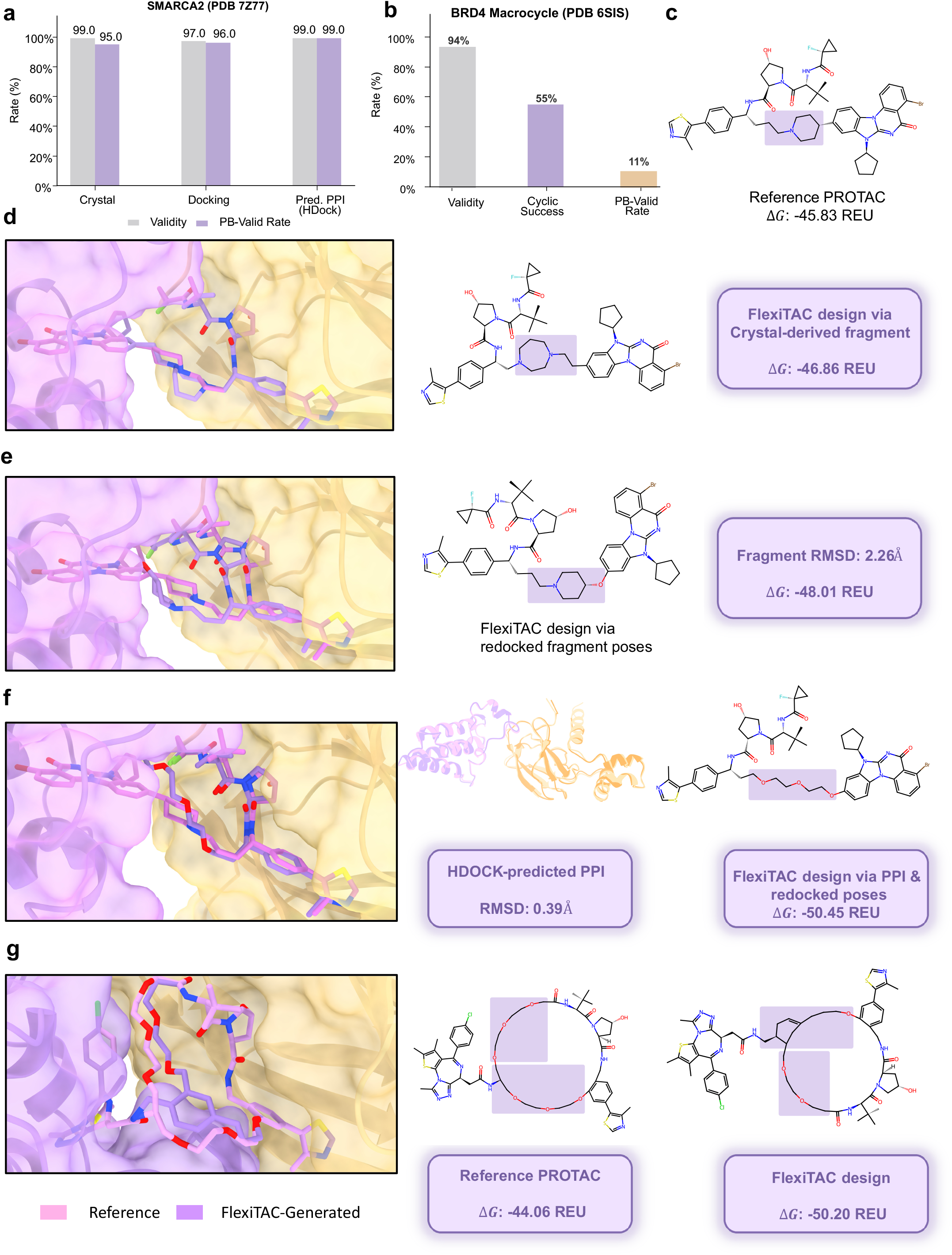
| PROTAC linker design across diverse scenarios with FlexiTAC. **a,** Validity and PoseBusters linker pass rates (PB-valid) for FlexiTAC-generated PROTAC linkers across three scenarios for PDB ID 7Z77. Grey bars indicate the fraction of chemically valid molecules; purple bars indicate the fraction passing all PoseBusters structural quality checks on the linker structures. **b,** Corresponding success rates for the design results for PDB ID 6SIS, showing the fraction of generated linkers that form a cyclic structure (Cyclic Success) alongside validity and PB-valid rate. **c,** Reference PROTAC from the 7Z77 crystal structure (ΔG: -45.83 Rosetta Energy Unit (REU)). **d,** FlexiTAC design via crystal-derived fragment (ΔG: -46.86 REU). **e,** FlexiTAC design via redocked fragment poses (fragment RMSD: 2.26 Å; ΔG, -48.01 REU). f, FlexiTAC design via HDOCK-predicted protein-protein interaction (PPI) complex and redocked fragment poses (PPI RMSD: 0.39 Å; ΔG, -50.45 REU). **g,** Macrocyclic PROTAC design results (PDB 6SIS) using crystal fragment poses, demonstrating generalization to cyclic linker topologies (reference ΔG: -44.06 REU; generated ΔG = -50.20 REU). In all panels, pink represents the structures of reference PROTACs, and purple represents the structures of FlexiTAC-generated PROTACs.

We generated 100 PROTACs for each fragment-pair systems, and evaluate the generated quality using validity and PoseBusters metrics, and estimated the binding-free energy of the PROTAC’s ternary complex using the PRosettaC^32,33^ protocol. For the fragment pair system from PDB 7Z77, FlexiTAC maintained high performance across all three settings (**Fig. 8a**). The validity rates were 99%, 97% and 99% for crystal-derived, redocked and predicted-complex inputs, respectively. The corresponding PB-valid rates were 95%, 96% and 99%. **Fig. 8c** shows the reference PROTAC and its linker topology.

Using the crystal-derived fragment poses, FlexiTAC generated a PROTAC that closely matched the ternary binding geometry, and finally obtained 14 PROTACs with better ΔG values than reference PROTAC. For the case shown in **Fig. 8d**, ΔG of the PROTAC’s ternary complex was -46.86 Rosetta Energy Unit (REU), compared with - 45.83 REU for the reference PROTAC. When the fragments were redocked using AutoDock-Vina^34^, FlexiTAC remained able to connect the two ligands despite a fragment RMSD of 2.26 Å (**Fig. 8e**). The representative design had an estimated ΔG of -48.01 REU, and obtained 8 PROTACs with better ΔG among the 100 generated PROTACs. FlexiTAC also generated a compatible PROTAC from an HDOCK^35^- predicted protein complex followed by fragment docking (**Fig. 8f**). The predicted protein complex had an RMSD of 0.39 Å, and resulted in 31 PROTACs with ΔG better than reference. The selected design had an estimated ΔG of -50.45 REU. Their spatial overlap and binding mode indicate that FlexiTAC can accommodate uncertainty of fragment pair condition introduced by docking and the predicted protein-protein complex.

We repeated the three-scenario experiment using the BRD4 PROTAC ternary complex from PDB 6BOY (**Supplementary Fig. 11**). The bar chart shows validity rates of 96%, 90% and 93% for the crystal, redocking and predicted-complex settings, respectively. The corresponding PB-valid rates were 96%, 90% and 90%, and yielded 71,60 and 59 PROTACs with better PRosettaC-estimated ΔG. The three structural panels show representative designs obtained from each input setting. The crystal-based design case had an estimated ΔG of -49.73 REU, compared with -49.34 REU for the reference PROTAC. The redocking-based design had a fragment RMSD of 0.24 Å and an estimated ΔG of -54.78 REU. The design based on the HDOCK-predicted complex had a protein RMSD of 0.35 Å and an estimated ΔG of -53.23 REU.

Linker design tasks predominantly focus on linear topological structures, which inherently limits model’s application in molecular generation task in more atypical chemical space. We sought to investigate whether FlexiTAC possesses the geometric adaptability and zero-shot generalization capacity to overcome this limitation and directly generate non-standard linker. Specifically, we examined the BRD4 PROTAC from PDB 6SIS (**Fig. 8g**), a PROTAC featuring a highly complex macrocyclic linker that requires more than two fragment-linker anchor points. We used FlexiTAC to generate 1,000 linkers for the fragment pair of 6SIS, and obtained chemically valid molecules in 94% of the samples, and 55% successfully formed a cyclic linker topology (**Fig. 8b**). The PB-valid rate was 11%, highlighting the greater geometric difficulty of cyclic linker generation. Nevertheless, the representative generated structure recovered the overall macrocyclic geometry of the reference PROTAC (**Fig. 8g**). Among the PROTACs that passed validity, cyclic and Posebusters filters, 18 PROTACs achieved better ternary complex energy than the original PROTACs in 6SIS. The selected case in **Fig. 8g** had a ΔG of -50.20 REU, compared with -44.06 REU for the reference PROTAC.

**Fig. 8** and **Supplementary Fig. 11** show that FlexiTAC can generate valid and structurally plausible PROTACs from crystal structures, redocked fragments and predicted protein complexes. The model also supports macrocyclic linkers with atypical anchor topologies. Although the PRosettaC values are computational estimates, they suggest that FlexiTAC-designed PROTACs can form stable ternary complexes and provide promising starting points for developing effective target degraders.

## Discussion

Successful PROTAC development depends on identifying linkers that reconcile productive ternary-complex formation with the physicochemical properties required for cellular activity. This challenge demands the joint optimization of chemical identity and 3D geometry under fixed fragment constraints. FlexiTAC addresses this problem using a BFN that jointly generates linker atom types and coordinates while preserving the input fragments. It achieves a strong balance between chemical validity and geometric quality across diverse input conditions, accommodating fragment poses from experimental structures, molecular redocking and predicted POI**-**E3 complexes, supporting partially specified inputs and enabling linker flexibility to be tuned through posterior-gradient guidance without retraining. By integrating discrete-continuous generation, heterogeneous structural inputs, incomplete-condition inference and conformational control within a unified framework, FlexiTAC advances linker design from empirical enumeration towards structure-aware and hypothesis-driven exploration. These capabilities provide a versatile platform for exploring linker chemistry and conformational space across different stages of PROTAC discovery and optimization

Beyond FlexiTAC, PROTAC-3D offers standardized, training-ready examples at a scale substantially exceeding existing public resources, whereas PROTAC-Bench enables consistent evaluation of fragment preservation, generalization, geometric fidelity, conformational stability, condition awareness and sampling efficiency. Together, these resources support the training, comparison and systematic diagnosis of future linker-generation models, making methodological progress more reproducible and directly comparable.

Together, FlexiTAC, PROTAC-3D and PROTAC-Bench establish an integrated model-data-benchmark paradigm for controllable linker design. By aligning structure-conditioned generation with task-specific data and application-relevant evaluation, this framework advances PROTAC linker exploration from empirical enumeration towards systematic, hypothesis-driven design. Although developed for PROTACs, the underlying formulation may inform other heterobifunctional and proximity-inducing modalities in which molecular components must be connected under geometric and physicochemical constraints. By bridging structural biology, generative modeling and medicinal chemistry, this work could ultimately expand the range of disease-relevant proteins and target-E3-ligase combinations amenable to targeted degradation, supporting the development of new therapeutic strategies.

## Acknowledgements

This work was supported by National Natural Science Foundation of China (82373937 to J.Y., U25A20568 to F.K.), Fundamental Research Funds for the Central Universities (22120260486) and Bioinformatics Supercomputing Center at the School of Life Sciences and Technology, Tongji University, for providing computational resources and support.

## Author Contributions Statement

Y.L., Y.Z., L.Z., C.H. and Q.X. contributed equally to this work. D.C. and J.Y conceived and supervised the research project. Y.L. developed the primary method and code. Y.L, Y.Z. and Q.X assisted in the analysis of the primary baselines and data. Y.L., Y.Z. and D.C. wrote the paper. All authors review and approved the final manuscript.

## Methods

### Pretraining Data Curation

Before PROTAC-3D training, we constructed a quasi-PROTAC dataset to expose FlexiTAC to diverse chemical structures and quantum-mechanically informed 3D conformations. We obtained 1.8 million molecules from ChEMBL-3D, whose conformations had been optimized using AIMNet2^36^.

Each molecule was represented as a molecular graph *M* = (*V*, *E*), where *V* represents atoms, and *E* represents chemical bonds. Two non-aromatic bonds, *b*_1_, *b*_2_ ∈ *E*, were randomly selected and cleaved to produce three connected components, (*F*_1_, *L*, *F*_2_). The resulting components *F*_1_ and *F*_2_ served as the conditioning fragments, whereas the intervening component *L* defined the target linker. An example was retained only when all three components contained at least three heavy atoms and could be parsed by RDKit. This procedure yielded 1,000,450 quasi-PROTACs.

The resulting dataset preserved AIMNet2-optimized geometries within a fragment-linker-fragment representation that is similar to the conditional structure of linker design. Its scale and chemical diversity were intended to improve conformational accuracy and broaden chemical coverage before fine-tuning on PROTAC-3D. Correspondingly, FlexiTAC (ChEMBL) achieved better JSD performance and relaxation stability compared to FlexiTAC (scratch).

### Bayesian Flow Network for Linker Generation

FlexiTAC formulates PROTAC design as a conditional linker generation problem. Given fixed E3-ligase-ligand and warhead fragments, the model generates the atoms and 3D coordinates of the linker that reconnects them into a complete PROTAC. Fragment atom types and coordinates remain conditioning information throughout training and sampling. The learning objective is the conditional distribution of linkers under its fragment-pair context.

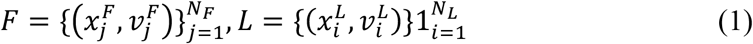

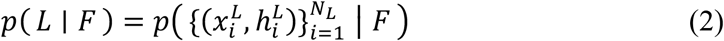

Here, *F* denotes the fragment-pair condition and *L* denotes the linker. *x^F^* and *x^L^* denote the fragment and linker coordinates, whereas *v^F^* and *v^L^* denote their atom types. *N_F_* and *N_L_* denote the corresponding atom counts

FlexiTAC is built on a BFN. Instead of adding noise directly to sampled molecules, the BFN maintains a distributional parameter for the linker. For coordinates, this state is represented by Gaussian parameters. For atom types, it is represented by categorical probabilities. The two modalities are therefore updated in a common parameter space, which avoids switching repeatedly between a noisy sample space and a discrete molecular graph as in diffusion models. We denote the parameter state at time *t* as:

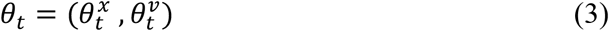

At each time step, the sender distribution *p_S_* produces a noisy observation of the true linker *L*. The SE(3)-equivariant backbone ϕ_ψ_ takes the current linker parameters *θ_t_*_-1_, the fragment condition and the time embedding *t* as input, and predicts the clean linker *L̂*, forming the BFN’s output distribution *p_0_*. The receiver distribution *p_R_* then takes the clean linker prediction of the *p_0_* and adds noise to it at the same noise level as the sender distribution. α_*i*_ denotes the noise schedule that controls the uncertainty degree of the sender and receiver distribution. A Bayesian update then moves the parameter state toward the data when the receiver agrees with the sender. In this way, the model learns a denoising direction in parameter space while preserving the geometric condition of the fragment pair.

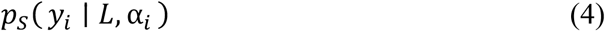

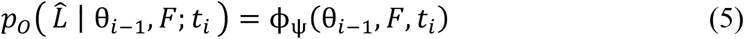

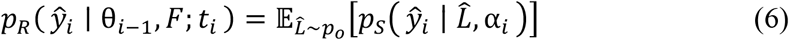

The coordinate uses continuous Bayesian Flow controlled by a coordinate noise scheduler, and the atom-type uses a continuous relaxation of categorical states controlled by an atom-type noise scale. Let *σ*_1_ denote the coordinate noise scale when BFN update, and γ(*t*) is the scheduler that controls the level of noise that is used in the sender and receiver distribution. Given the true linker coordinates *x_L_*, the Bayesian flow distribution over the atom coordinates is sampled as:

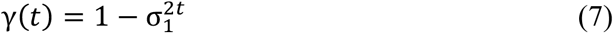

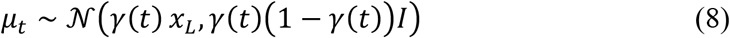

In the context of the BFN parameter update, 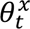 consists of 2 parts {*µ_i_*, *ρ_i_*}, where the *ρ_i_* is a predefined schedule that controls how much information is revealed at each timestep, and *µ_i_* is the mean of the Gaussian distribution, the closed-form Bayesian update for coordinates is:

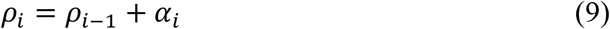

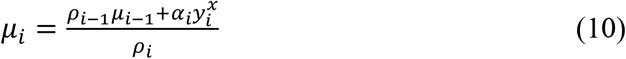

For discrete atom types with K categories, FlexiTAC uses softmax logits for parameterization. Let ***e***(ℎ*^L^*) denote the one-hot vector of the true atom type and let *β* denote the noise scale for atom type. the discrete Bayesian flow for atom type is defined as:

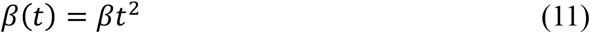

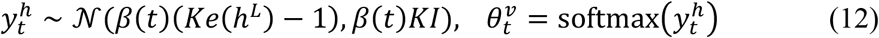

During training, a discrete time index *i* is sampled and the linker parameter state is constructed. The coordinate loss and the atom-type loss measure the KL divergence between predicted and true linker. The final BFN loss is the weighted sum of these two terms. In our implementation, the atom-type term is up-weighted by *λ*_type_to balance with coordinate loss.

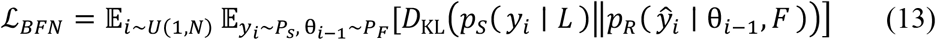

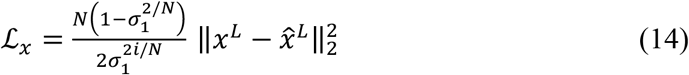

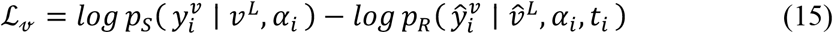

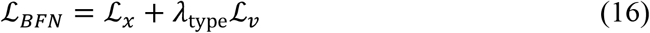

During sampling, the linker starts from an uncertain parameter state. FlexiTAC repeatedly predicts a clean linker estimate and applies the Bayesian update, gradually reducing uncertainty in both coordinates and atom types. After the final step, the predicted linker is reconstructed together with the two unchanged fragments. Fragment topology is preserved from input molecules, and linker-fragment bonds are assigned through the predefined or predicted anchor atoms.

### Posterior Guidance for Flexibility Control

To enable explicit control over linker flexibility, we defined a conformational ensemble-derived rigidity score (CERScore), denoted *r*_rigid_. CERScore integrates spatial atom distance, anchor-distance stability and global intralink shape stability. For each PROTAC, we generated 100 conformers using RDKit and optimized them with the MMFF94 force field^30^. Conformational fluctuations were evaluated across this force-field-minimized ensemble. Larger fluctuations indicate lower geometric stability and therefore produce lower rigidity scores.

Local topological freedom was quantified using the number of rotatable bonds in the linker:

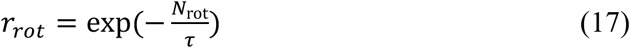

where *N*_rot_ is the number of linker rotatable bonds. The constant *τ* = 5.0 controls the decay rate and smooths the resulting score distribution.

Anchor-distance stability was measured from fluctuations in the distance between the two linker attachment atoms:

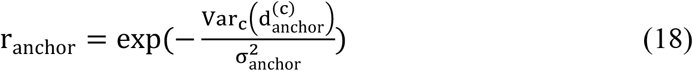

where 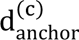 is the anchor-to-anchor distance in conformer *c*. The term 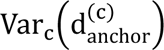 denotes its Boltzmann-weighted variance across the conformational ensemble.

Global shape stability was quantified using fluctuations in long-range intralink distances:

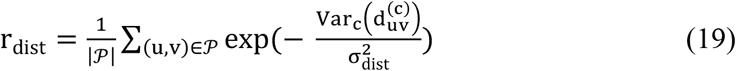

where *P* contains all atom pairs separated by at least three topological bonds along the linker. For each pair (*u*, *v*), 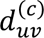 is their Euclidean distance in conformer *c*. Its Boltzmann-weighted variance measures the stability of the corresponding long-range intralink geometry.

The final CERScore was calculated as a weighted combination of the three terms:

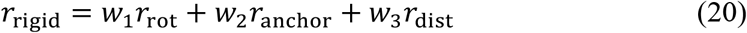

The resulting score satisfies *r*_rigid_ ∈ [0,1], with higher values indicating greater rigidity. Flexible linkers contain more rotatable bonds and exhibit larger conformational fluctuations, thereby receiving lower rigidity scores.

A rigidity prediction head *ϕ_rigid_* is trained to provide a differentiable estimate of CERScore. The head takes hidden linker embeddings from the FlexiTAC backbone and maps them to a value between 0 and 1. The backbone is frozen and only the prediction head is optimized against the precomputed rigidity labels when training, we use smooth L1 loss to calculate the prediction loss:

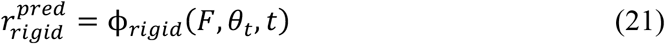

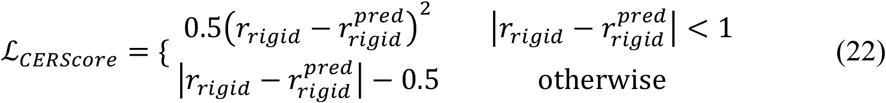

During guided generation, FlexiTAC uses the gradient of the CERScore prediction head ϕ*_rigid_* with respect to the current linker state. The coordinate prediction and the atom-type logits are shifted along the coordinates and atom-type gradient before the next Bayesian update. Guidance has been applied during the last 60 sampling steps,and the range of *t* is 40-100 when the number of BFN inference steps is 100. Positive guidance weights bias generation towards more rigid linkers, whereas reversing the sign favors more flexible linkers. The hyperparameters *λ_x_* and λ*_v_* denote the guidance intensity.

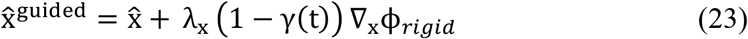

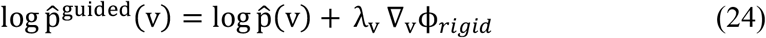

Where xN and log pN(v) are the original linker atom coordinates and the log-probability of atom types predicted by FlexiTAC, ∇_x_ϕ*_rigid_* and ∇_v_ϕ*_rigid_* are the rigidity prediction head’s gradient calculated for coordinates and atom-type. The hyperparameters *λ_x_* and λ*_v_* denote the guidance intensity. γ(t) is the coordinate noise level at time t, the factor (1 − γ(t)@) suppresses guidance intensity at the earlier high noise steps, and uses stronger guidance intensity at later low noise steps to provide a clear guiding inference trajectory.

### Auxiliary Modules

FlexiTAC can use the reference linker size or predict the linker size before BFN sampling. The linker size prediction module is an Equivariant Graph Neural Network (EGNN) that uses fragment-pair geometry, to estimate the number of linker heavy atoms. This module allows generation when the desired linker length is not known.

The anchor prediction module estimates which fragment atoms should connect to the linker. It is also an EGNN network that takes fragment pair as input. Both modules are trained on the same training data of PROTAC-3D as the FlexiTAC backbone model. Together, the size and anchor modules reduce the amount of manual information needed for generation, while keeping the core linker generator process unchanged.

The Size GNN is trained using a Smooth L1 loss to predict the number of linker atoms, where *B* is the batch size, *ŷ_b_* and *y_b_* are the predicted and ground-truth number of linker atoms for sample *b*, respectively, and *β* is the transition parameter, *δ* = *ŷ_b_* − *y_b_*:

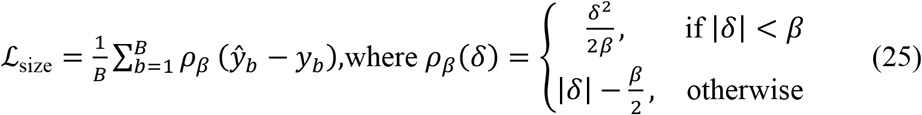

The Anchor Prediction Head is trained using a weighted Binary Cross-Entropy (B CE) loss, where 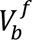 is the set of fragment atoms in sample *b*, *p_bi_* is the predicted prob ability of atom *i* being an anchor, *y_b_i__* ∈ {0,1} is the ground-truth anchor label:

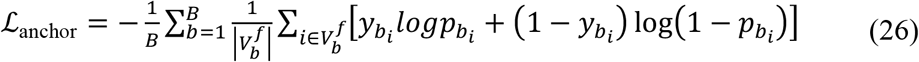

### Training Details

The FlexiTAC backbone uses an SE(3)-equivariant graph neural network with 9 layers, a hidden dimension of 128, 16 attention heads and 20 radial Gaussian features. Neighborhoods are constructed with k-nearest-neighborhood (KNN) graph of 32 nearest neighbors, and the coordinate cutoff radius is 10 Å.

FlexiTAC is trained with AdamW, a learning rate of 1e-4, weight decay of 1e-8 and exponential moving average decay of 0.999. The default batch size is 64. The model is trained on PROTAC-3D for 90 epochs, with an atom-type loss weight of 10.0. A plateau scheduler reduces the learning rate by a factor of 0.6 after five epochs without a decrease in validation loss, with a minimum learning rate of 1e-6. The FlexiTAC ChEMBL-pretrain version was first trained on our quasi-PROTAC dataset for 35 epochs, then trained on PROTAC-3D for the same 90 epochs.

During evaluation, FlexiTAC uses 100 sampling steps and generates 100 candidate linkers for each fragment pair in PROTAC-Bench curated test sets. The linker-size prediction module is trained as a regression model with a 4-layer graph network and Smooth L1 loss on the PROTAC-3D dataset for 100 epochs. The anchor prediction module uses the same network architecture as the linker-size prediction module and is trained as a classification network. The training target is to assign the highest probability for the anchor atom on each fragment and is conducted on the PROTAC-3D dataset for 100 epochs.

### PROTAC-Bench construction and evaluation

Initially, all the terminal fragment pairs in PROTAC-3D dataset were canonicalized before system grouping and split assignment. We defined each unique warhead-E3-ligand pair as one fragment-pair system, which serves as the conditional information for the model. All PROTACs containing the same pair served as reference PROTAC molecules for that fragment-pair system. A component was considered unseen if it was absent from the training data. Both-unseen systems contained an unseen warhead and an unseen E3 ligand. E3-unseen systems paired an unseen E3 ligand with a seen warhead, whereas warhead-unseen systems used the converse configuration. The benchmark comprised 30 both-unseen, 50 E3-unseen and 50 warhead-unseen systems. The 130 systems comprised 3,516 reference PROTACs in total: 555 in both-unseen, 1,083 in E3-unseen and 1,878 in warhead-unseen. Reference counts per system ranged from 10 to 27 in both-unseen (median, 19). They ranged from 2 to 121 in E3-unseen (median, 14.5) and 20 to 105 in warhead-unseen (median, 31.5). Notably, many E3-unseen ligands retained a cereblon (CRBN)-binding core because CRBN recruiters predominate in patent-derived PROTAC data. These ligands were nevertheless structurally distinct derivatives. Even small substitutions can alter ternary-complex stability and the geometry presented to the linker. Retaining these variants therefore tests robustness to subtle, functionally relevant changes within the dominant E3-ligase class.

Metrics were calculated first for individual generated samples, then summarized within fragment-pair systems and finally across each test setting. A sample was valid only if RDKit sanitization succeeded, the molecule was connected, both input fragments were preserved, and a standardized linker could be extracted. Validity was calculated over all requested samples. Subsequent molecular descriptors and reference comparisons used valid samples, unless a metric explicitly measured generation failure. Metrics were first evaluated on PROTACs from the same fragment-pair system. Then, fragment-pair level evaluation results were averaged within each test set (E3-unseen, warhead-unseen and both-unseen). This hierarchy prevented reference-rich systems from dominating the benchmark.

Hit rate asks whether a model-generated PROTACs match or resemble any reference PROTACs of its input fragment pairs. Strict hits require canonical SMILES identity. Similarity-based hits used the whole-PROTAC Morgan Tanimoto similarity and were evaluated at 1.0, >0.9, >0.8. Hit rate was the number of hits within the generated PROTACs of each model. Hit fraction quantified reference-set coverage by asking how many reference PROTACs could be recovered. Each reference was counted once when any valid generated sample reached the selected similarity threshold. Hit fraction was the number of recovered references divided by the total unique reference count for that system. Hit rate and hit fraction therefore provide complementary views of precision and coverage, enabling a more complete assessment of conditional generation.

We also reported Shape and Color similarity score(SC-RDKit), which combines shape overlap and pharmacophore agreement with a reference conformer. Generated conformers with SC-RDKit scores above 0.95 were counted as passing. Conformational robustness was assessed by the structural change required during force-field relaxation. MMFF94 energies were recorded before and after minimization, and their difference was reported as Δenergy. We also measured minimization-induced linker RMSD, conformational strain energy and PoseBusters^37^ pass rates. These metrics distinguish a chemically valid graph from a conformation that remains plausible after local relaxation. Practical PROTAC design spans ternary geometries in which the two linker attachment atoms can be either close together or widely separated. We therefore designed an anchor-distance test to assess whether 3D linker generators remained valid across this range. The analysis included 90 fragment pair systems spanning anchor distance of 3.108-27.057 Å on 43 both-unseen, 25 E3-unseen and 22 warhead-unseen. Five 3D models, FlexiTAC (ChEMBL), FlexiTAC (scratch), PocketXMol, LinkerNet and DiffLinker, each generated 100 candidates per fragment pair system. Systems were assigned to nine regions from anchor distance close to far, with ten fragment-pair systems per region. We reported the mean validity of the generated PROTACs within each region.

A linker-generation model should use the chemical information encoded by its input fragments rather than merely construct linker-like structures between two attachment points. We therefore developed the Fragment-Aware Score (FAScore) to measure whether a model preferentially recovers the appropriate reference linkers when conditioned on their corresponding fragment pair.

For model (m) and fragment pair (i), FAScore was defined as

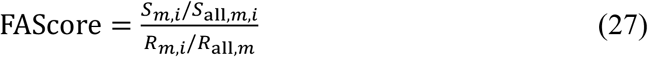

Where *S*_all,*m*,*i*_ is the number of distinct valid linkers generated by model m under fragment condition i; *S_m,i_* is the number of these conditionally generated linkers that recover at least one reference linker associated with fragment pair i; *R*_all,*m*_ is the number of distinct valid linkers generated by the same model across all 130 fragment pairs; and *R_m,i_* is the number of linkers in this global background that recover on of the reference-linkers for fragment pair i.

A FAScore greater than 1 indicates that the correct fragment condition enriches its associated reference linkers above the model’s global background. A score of 10 indicates tenfold enrichment. Fragment pairs without conditional recovery were assigned a score of zero. FAScores above 100 were retained in the numerical results but capped at 100 for visualization. **Fig. 3** reports the fraction of fragment pairs with FAScore <10 and with 10 ≤ FAScore ≤ 100.

To evaluate the model’s inference-time scaling ability as a practical tool for scalable PROTAC discovery, we selected one reference-rich fragment system from each test set of PROTAC-Bench. Their reference PROTACs contained 16 both-unseen, 121 E3-unseen and 88 warhead-unseen PROTACs. FlexiTAC, DiffLinker and LinkerNet were run for up to 1.5 h per system, and reference recovery was evaluated at 300-s intervals. Recovery was assessed using whole-PROTAC Morgan Tanimoto similarity thresholds of 0.8. At each time point, we reported hit fraction, which quantifies the proportion of unique references hit, together with the corresponding hit count. The resulting time-based curves measure how rapidly each model covers its reference PROTAC set under the same inference time. Similarly, we tracked the statistics based on the total number of samples. For every 100 PROTACs sampled, we calculated the hit rate and hit count for each model to plot a sample-size-based curve.

Hit rate and hit fraction are evaluated based on the recovery of the whole PROTAC, which does not capture motif-level coverage. We therefore screened 3,516 valid reference linkers from all reference PROTACs in the test sets against a curated library of SMARTS patterns of common functional groups and ring scaffolds. Ring systems occurring in at least three linkers and not duplicating a curated motif were then added. After retaining only motifs that occur in at least one real test linker, the final reference panel comprised 62 reference motifs. Generated linkers were canonicalized and deduplicated before substructure matching evaluation. A motif was counted as covered when at least one unique generated linker matched its pattern.

### Baselines

We compared FlexiTAC with seven representative baselines, organized into two broad families according to whether they generate linkers in two-dimensional molecular space or explicitly model 3D fragment geometry. The 2D baselines were DeLinker and Link-INVENT^38^. DeLinker represents a graph-based fragment-linking model conditioned on two molecular fragments and their attachment geometry, whereas Link-INVENT extends the REINVENT^39^ framework to linker generation with a sequence-based generative policy. DeLinker was retrained on the PROTAC-3D training split, while Link-INVENT was evaluated with its released pretrained checkpoint through the REINVENT4 sampling workflow.

The 3D baselines include 3DLinker^40^, two equivariant diffusion models, and transferable 3D molecular generation frameworks. 3DLinker was included as a linker-specific 3D generative model that jointly predicts linker topology and coordinates from fixed fragment poses. DiffLinker and LinkerNet were evaluated as representative equivariant diffusion baselines: DiffLinker denoises linker atom types and coordinates under fragment constraints, whereas LinkerNet performs fragment-pose and linker co-design with an equivariant diffusion model. Both diffusion models were adapted to the fixed-fragment PROTAC setting and retrained on PROTAC-3D. We further evaluated FFLOM^41^ and PocketXMol^42^ as transferable 3D generation baselines. FFLOM was retrained on processed PROTAC-3D tensors and molecule SMILES, whereas PocketXMol was used directly with its released pretrained checkpoint and constrained at sampling time to preserve the two PROTAC fragments.

We also trained FlexiTAC on PROTAC-DB 1.0 and ZINC training datasets and used the same test sets to evaluate FlexiTAC’s performance to provide a fair comparison. Detailed information and evaluation results are provided in the **Supplementary Information.**

For controlled evaluation, all retrainable baselines were trained with their original objectives and codebases on the same PROTAC-3D training split. Thus, DeLinker, 3DLinker, DiffLinker, LinkerNet and FFLOM were retrained, while Link-INVENT and PocketXMol were evaluated by direct inference from official pretrained weights. Because these baselines require task-specific oracle information, we supplied the reference linker size and anchor-atom labels during sampling when required by the model interface. Each method generated 100 candidate linkers for every fragment pair in both-unseen, E3-unseen and warhead-unseen test sets. Full implementation details, checkpoint sources and conditioning settings are provided in the **Supplementary Information.**

## Data Availability

All the processed datasets, as well as pretrained models, are available at zenodo a nd Github. The datasets that used by the baselines are the ZINC dataset(https://doi.org/10.5281/zenodo.7121271); and PROTAC-DB 1.0 (https://github.com/guanjq/LinkerNet). PROTAC-3D and PROTAC-Bench’s test sets are accessible at https://doi.org/10.5281/zenodo.22030032. Crystal structures in case studies are available at Protein Data Bank under the accession codes 6BOY, 7Z77 and 6SIS, respectively.

## Code Availability

The code used to generate the results reported in this study will be made publicly available upon acceptance at the following GitHub repository: https://github.com/Intelligent-Drug-Discovery-Lab/FlexiTAC.

