## Supplementary Information for "FlexiTAC enables controllable PROTAC linker generation across diverse structural settings using a Bayesian flow network with posterior guidance"

**Table of Contents**

**S1 Supplementary Information for PROTAC-3D dataset**

**S2 Supplementary Information for the linker size and anchor atom prediction modules**

**S3 Supplementary Information for the PROTAC-DB 1.0 and ZINC evaluation results**

**S4 Supplementary Information for Jensen-Shannon divergence distribution s of linker substructures**

**S5 Supplementary Information for motif coverage**

**S6 Supplementary Information for the rigidity guidance sampling results**

**S7 Supplementary Information for the multi-scenario application of FlexiTAC**

**S8 Supplementary Information for baseline models on PROTAC-Bench**

**S9 References**

### S1 Supplementary Information for PROTAC-3D dataset

We first compared the training usability and structural diversity of PROTAC-3D with those of PROTAC-DB 3.0. PROTAC-DB 3.0 contained 9,380 source records, of which 6,110 remained after canonicalization. Requiring a complete tripartite decomposition retained 4,873 molecules, and subsequent structural quality control yielded 4,523 model-usable molecules. Cluster normalization further reduced this set to 2,849 representative triplets. By comparison, all 63,554 PROTAC-3D records remained canonicalizable, contained a complete warhead-linker-E3-ligand decomposition and passed the model-usability criteria. Cluster normalization at a generalized Tanimoto threshold of 0.7 retained 51,755 training-ready triplets (**Supplementary Fig. 1a**). To avoid inflating component diversity through minor structural variations, warheads and E3 ligands were grouped using Morgan count fingerprints at the same threshold. PROTAC-3D contained 16,563 canonical linkers, 5,075 warhead clusters and 2,570 E3-ligand clusters, compared with 1,301, 326 and 49, respectively, in PROTAC-DB 3.0 (**Supplementary Fig. 1b**).

Nearest-neighbour analysis further distinguished the two datasets. Similarity distributions for both linkers and full PROTACs were shifted towards lower values in PROTAC-3D. By contrast, PROTAC-DB 3.0 showed a pronounced accumulation of structures with similarities approaching  (**Supplementary Figs. 1c,d**). This pattern is consistent with greater representation of duplicate, fingerprint-equivalent or closely related analogues in PROTAC-DB 3.0. Heatmaps of the 12 most frequent warhead and E3-ligand clusters further showed that PROTAC-3D associated distinct linkers with more fragment-pair combinations (**Supplementary Figs. 1e,f**). Although common E3 ligands dominated both datasets, PROTAC-3D provided broader linker coverage beyond the most frequent warhead-E3-ligand combinations.

The linker-property distributions occupied broadly overlapping physicochemical space but differed in polarity, flexibility and ring content (**Supplementary Fig. 2**). PROTAC-3D provided greater coverage of non-zero hydrogen-bond acceptor and donor count without markedly shifting the overall topological polar surface area or lipophilicity distributions. More pronounced differences were observed for rotatable bonds and ring-related descriptors. Linkers in PROTAC-3D had fewer rotatable bonds and more rings than those in PROTAC-DB 3.0, with median ring counts of 1 and 0, respectively. The median maximum ring size was six atoms in PROTAC-3D but zero in PROTAC-DB 3.0. PROTAC-DB 3.0 was therefore enriched in acyclic, conformationally flexible linkers, whereas PROTAC-3D provided greater representation of cyclic and heterocycle-containing architectures.

This distinction is relevant to linker design because flexibility and conformational constraint have complementary roles. Flexible linkers allow the two ligands to sample alternative relative orientations. However, excessive flexibility expands the accessible conformational ensemble and may reduce the probability of adopting a productive ternary-complex geometry. Cyclic linkers restrict this ensemble and can reorganize the linked ligands, although the optimal balance depends on the target, ligands and ternary-complex geometry. These distributions therefore indicate that PROTAC-3D covers a broader flexibility-constraint spectrum, rather than establishing that cyclic linkers are universally superior.

At the full-molecule level, PROTAC-3D also occupied broad PROTAC-relevant physicochemical space (**Supplementary Fig. 3**). Its distributions were generally smoother than those of PROTAC-DB 3.0 and shifted towards lower molecular weight, topological polar surface area, hydrogen-bond donor count and rotatable-bond count. Much of the distribution overlapped the literature-based reference regions for molecular weight, topological polar surface area, hydrogen-bond acceptors, hydrogen-bond donors and cLogP. Although the rotatable-bond distribution extended beyond the reference range, these values provide contextual benchmarks rather than strict cut-offs for beyond-rule-of-five PROTACs. Overall, the expanded structural coverage did not shift PROTAC-3D towards extreme physicochemical properties, although these descriptors alone cannot establish permeability or oral bioavailability.

The most recurrent linker structures provided a chemical explanation for these distributional differences (**Supplementary Fig. 4**). All ten top-ranked linkers in our dataset contained at least one ring, including piperidine, piperazine, cycloalkyl and aryl motifs. By contrast, the ten most frequent linkers in PROTAC-DB 3.0 were predominantly linear, acyclic aliphatic chains, some bearing a terminal carbonyl group. This enrichment of cyclic architecture is consistent with the increasing use of conformationally constrained linkers during PROTAC lead optimization and may reflect the greater representation of later-stage medicinal chemistry in patents. Such linkers restrict conformational sampling and can favor productive ternary-complex geometries, although conformational restriction alone does not determine degradation activity or permeability. Notably, the highest-ranked linker in our dataset corresponded to the piperidine-methylene-piperazine architecture used in Veppanu, which became the first FDA-approved PROTAC on 1 May 2026. This clinical precedent illustrates how patent-derived data can complement public databases with design motifs that have progressed through advanced drug development.

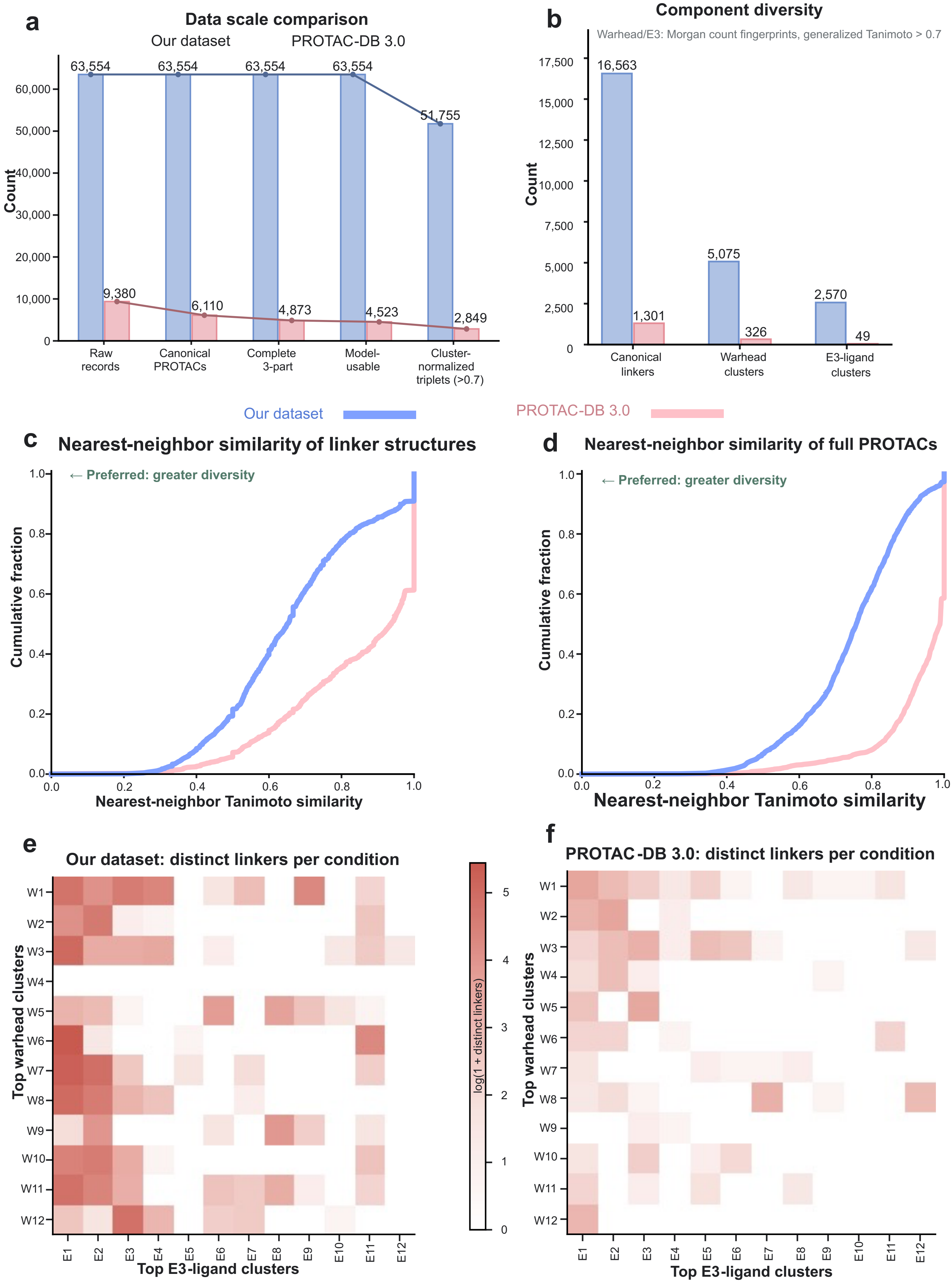
a
Data scale comparison
Our dataset
PROTAC-DB 3.0
Count
63,554
63,554
63,554
63,554
51,755
9,380
6,110
4,873
4,523
2,849
Raw records
Canonical PROTACs
Complete 3-part
Model-usable
Cluster-normalized triplets (>0.7)
b
Component diversity
Warhead/E3: Morgan count fingerprints, generalized Tanimoto > 0.7
16,563
1,301
5,075
326
2,570
49
Canonical linkers
Warhead clusters
E3-ligand clusters
Our dataset
PROTAC-DB 3.0
c
Nearest-neighbor similarity of linker structures
← Preferred: greater diversity
Cumulative fraction
Nearest-neighbor Tanimoto similarity
d
Nearest-neighbor similarity of full PROTACs
← Preferred: greater diversity
Cumulative fraction
Nearest-neighbor Tanimoto similarity
e
Our dataset: distinct linkers per condition
Top warhead clusters
Top E3-ligand clusters
log(1 + distinct linkers)
f
PROTAC-DB 3.0: distinct linkers per condition
Top warhead clusters
Top E3-ligand clusters

**Supplementary Fig. 1 | Dataset scale, component diversity and structural redundancy of the training data. a,** Sequential filtering of raw records into canonical PROTACs, molecules with complete three-part decomposition, model-usable records and cluster-normalized triplets. **b,** Numbers of canonical linkers and warhead and E3-ligand clusters, with fragment clusters defined using Morgan count fingerprints and a generalized Tanimoto similarity threshold of 0.7. **c,d,** Cumulative distributions of nearest-neighbor Tanimoto similarity for linker structures and full PROTACs, where lower similarity indicates lower structural redundancy. **e,f,** Numbers of distinct linkers associated with combinations of the 12 most abundant warhead and E3-ligand clusters in our dataset and PROTAC-DB 3.0; color intensity represents log(1 + number of distinct linkers). Blue denotes our dataset and orange denotes PROTAC-DB 3.0. Overall, our dataset contains more training-ready records, structurally distinct components and warhead-E3-ligand combinations while exhibiting lower nearest-neighbor similarity, indicating broader and less redundant chemical coverage.

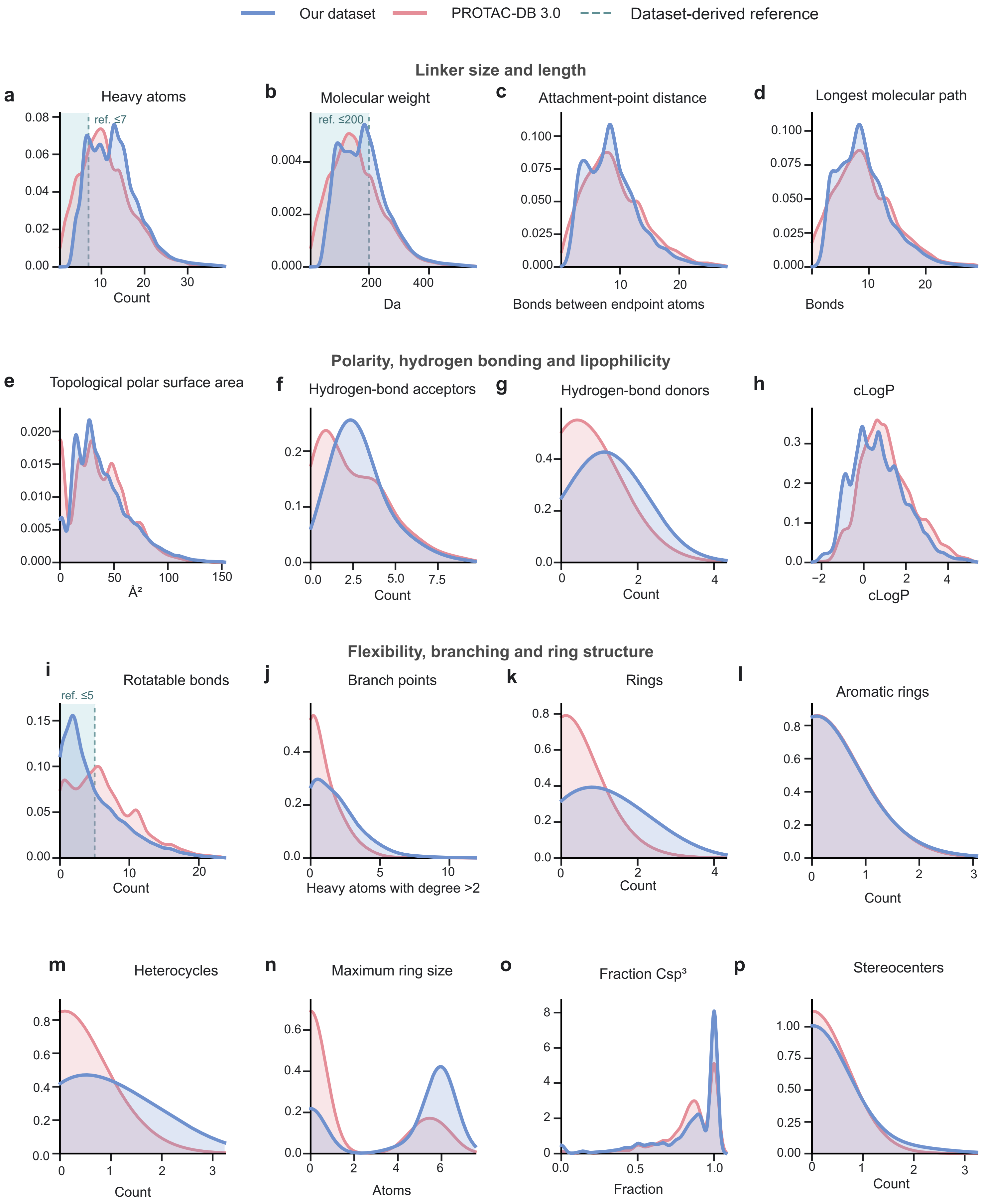

Our dataset
PROTAC-DB 3.0
Dataset-derived reference
Linker size and length
a
Heavy atoms
ref. ≤7
Count
b
Molecular weight
ref. ≤200
Da
c
Attachment-point distance
Bonds between endpoint atoms
d
Longest molecular path
Bonds
Polarity, hydrogen bonding and lipophilicity
e
Topological polar surface area
Å²
f
Hydrogen-bond acceptors
Count
g
Hydrogen-bond donors
Count
h
cLogP
cLogP
Flexibility, branching and ring structure
i
Rotatable bonds
ref. ≤5
Count
j
Branch points
Heavy atoms with degree >2
k
Rings
Count
l
Aromatic rings
Count
m
Heterocycles
Count
n
Maximum ring size
Atoms
o
Fraction Csp³
Fraction
p
Stereocenters
Count

**Supplementary Fig. 2 | Physicochemical and topological property distributions of extracted linkers.** Density plots compare linker size and length (**a-d**), polarity, hydrogen-bonding capacity and lipophilicity (**e-h**), and flexibility, branching and ring-related properties (**i-p**) between our dataset and PROTAC-DB 3.0. All distributions are normalized to unit area. Grey dashed lines and lightly shaded regions indicate empirical benchmarks for heavy-atom count (≤7), molecular weight (≤200 Da) and rotatable-bond count (≤5)[1,2]. The two datasets occupy overlapping core linker-property space, whereas the broader distributions observed for several sizes, hydrogen-bonding, branching and ring descriptors demonstrate that our dataset extends the structural diversity available for linker-generation model training.

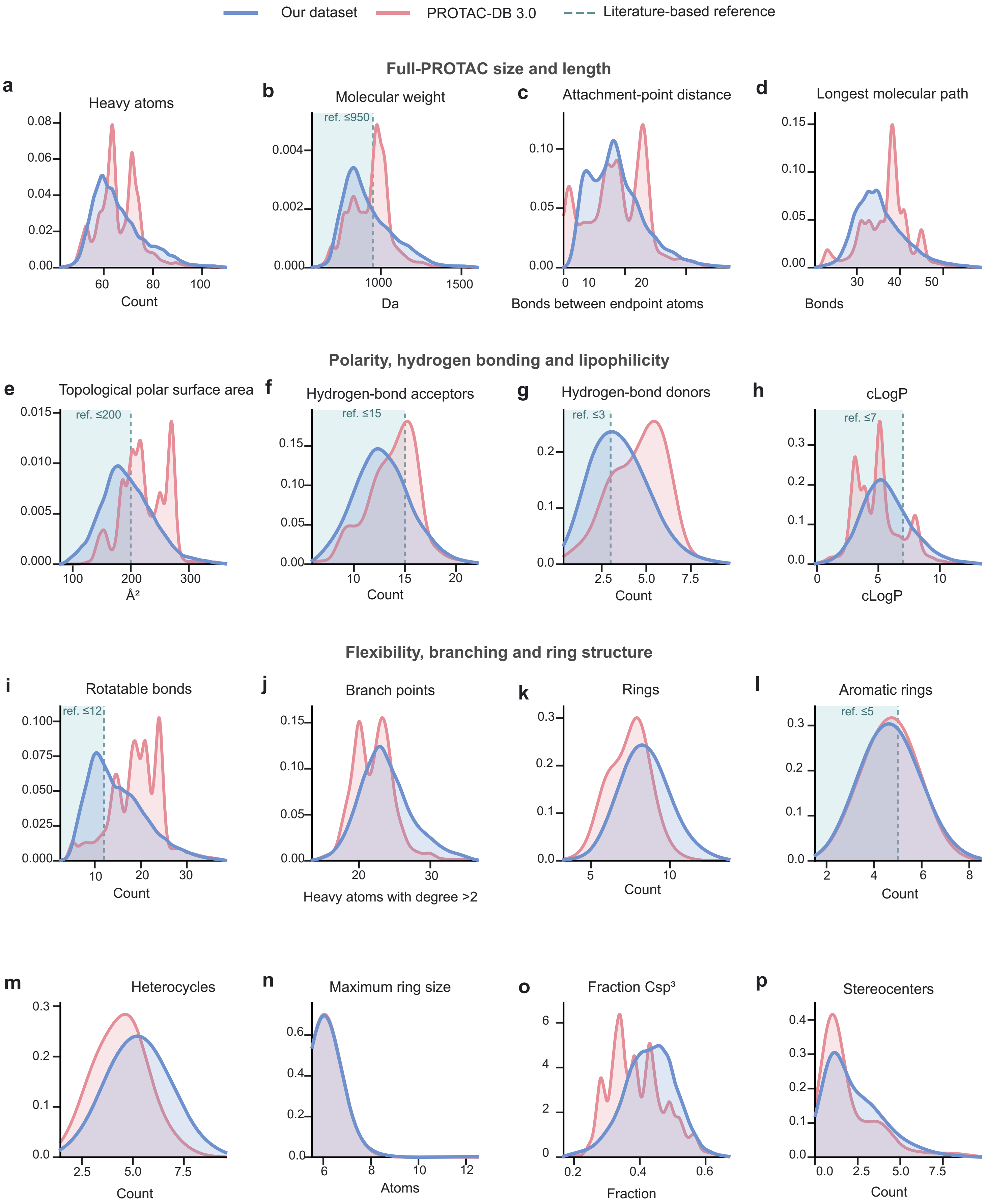
Our dataset
PROTAC-DB 3.0
Literature-based reference
Full-PROTAC size and length
a
Heavy atoms
Count
b
Molecular weight
ref. ≤950
Da
c
Attachment-point distance
Bonds between endpoint atoms
d
Longest molecular path
Bonds
Polarity, hydrogen bonding and lipophilicity
e
Topological polar surface area
ref. ≤200
Å²
f
Hydrogen-bond acceptors
ref. ≤15
Count
g
Hydrogen-bond donors
ref. ≤3
Count
h
cLogP
ref. ≤7
cLogP
Flexibility, branching and ring structure
i
Rotatable bonds
ref. ≤12
Count
j
Branch points
Heavy atoms with degree >2
k
Rings
Count
l
Aromatic rings
ref. ≤5
Count
m
Heterocycles
Count
n
Maximum ring size
Atoms
o
Fraction Csp³
Fraction
p
Stereocenters
Count

**Supplementary Fig. 3 | Physicochemical and topological property distributions of full PROTAC molecules.** Density plots compare full-PROTAC size and length (**a-d**), polarity, hydrogen-bonding capacity and lipophilicity (**e-h**), and flexibility, branching and ring-related properties (**i-p**) between our dataset and PROTAC-DB 3.0. All distributions are normalized to unit area. Grey dashed lines and lightly shaded regions denote literature-based reference values for molecular weight (≤950 Da), topological polar surface area (≤200 Å²), hydrogen-bond acceptors (≤15), hydrogen-bond donors (≤3), cLogP (≤7), rotatable bonds (≤12) and aromatic rings (≤5). Compared with PROTAC-DB 3.0, our dataset provides broader full-molecule coverage while showing greater representation in lower molecular-weight, polarity, hydrogen-bonding and flexibility ranges, supporting the physicochemical relevance and diversity of the training set.

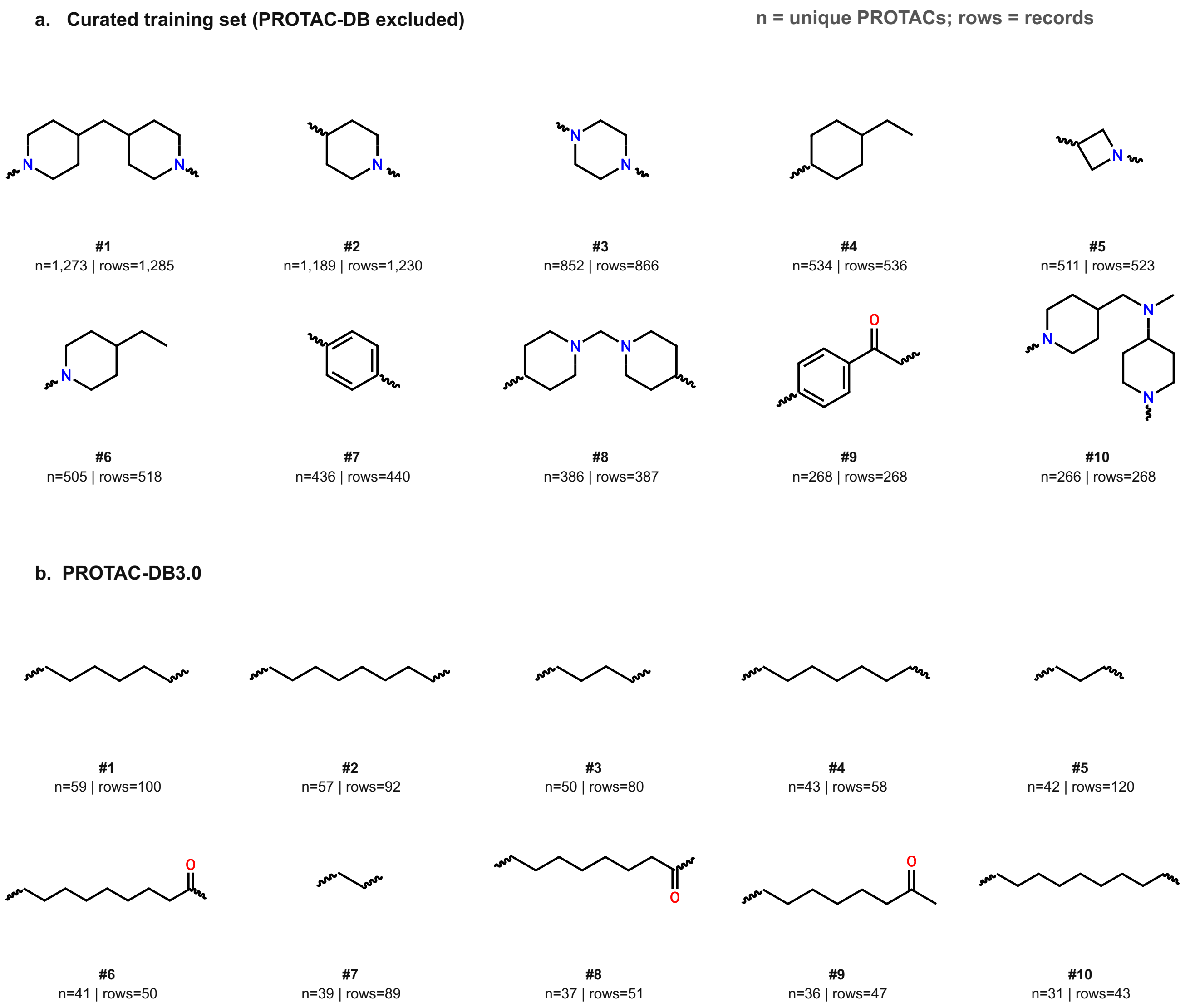


**Supplementary Fig. 4 | Most frequently occurring linker structures in the curated training set and PROTAC-DB**. The top 10 linker structures are ranked by their frequencies in our PROTAC-3D training set after removing PROTAC-DB entries and in PROTAC-DB itself. For each linker, “n” indicates the number of unique PROTACs containing that structure, whereas “rows” indicates the total number of associated records. The comparison highlights differences in recurrent linker motifs and their representation across the two datasets.

### S2 Supplementary Information for the linker size and anchor atom prediction modules

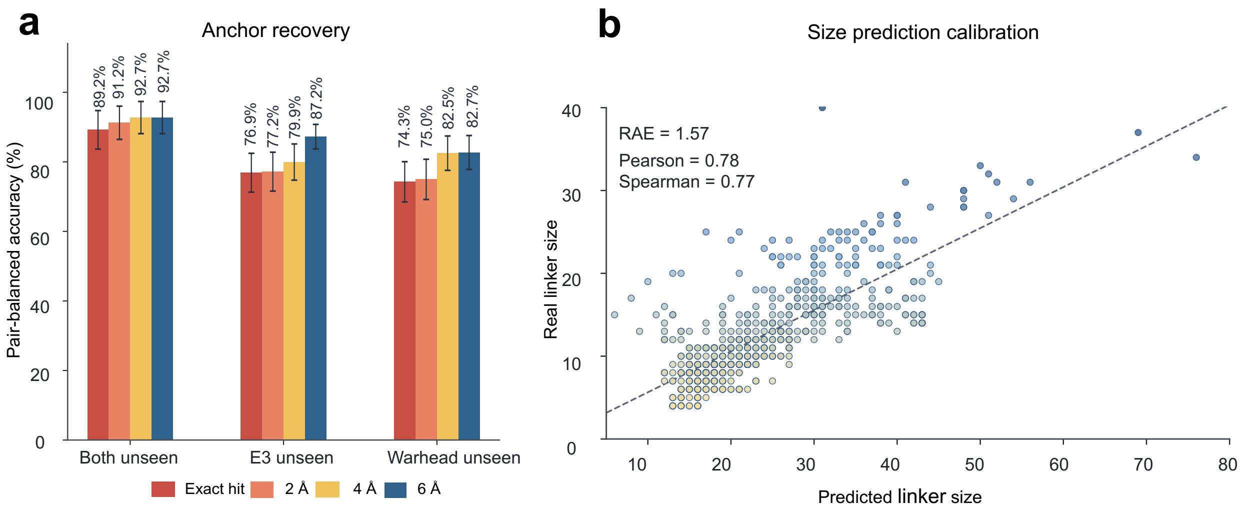


**Supplementary Fig. 5 | Performance of anchor and linker size prediction modules. a,** The anchor-atom prediction module was evaluated by whether the predicted anchor positions fell within different spatial tolerances of the true anchor atoms. The success rate increased from top-1 exact recovery to 2Å, 4 Å and 6 Å regions. **b,** The linker-size prediction module was assessed by comparing predicted linker size with the true linker size.

For the auxiliary prediction modules, we evaluated anchor localization and linker-size estimation on PROTAC-Bench test sets. For the anchor module, the model assigns an anchor probability to each fragment atom. We first measured the strict top-1 recovery, where the highest-scoring atom must exactly match a true linker-attachment atom. We then used a more tolerant region-based criterion: the predicted top-ranked anchor atoms were considered successful if they fell within 2 Å, 4 Å or 6 Å of the true anchor atoms in the fragment conformation. Under this evaluation, exact anchor recovery was 89.2%, 76.9% and 74.3% in both-unseen, E3-unseen and warhead-unseen test sets, respectively. The model performs best on the both-unseen set primarily because these molecules exhibit simpler geometric features, such as the shortest mean anchor distance of 11.3 Å and the lowest mean linker atom count of 10.8. These shorter distances and less complex structures facilitate more accurate anchor predictions, with the overall success rate further increasing within the 2 Å, 4 Å, and 6 Å regions.

For linker-size prediction, the same script compares the predicted linker atom count with the ground-truth linker size for each fragment pair. The calibration plot shows a clear positive association between predicted and true linker sizes, with Pearson r = 0.78 and Spearman r = 0.77. This suggests that the size module captures the monotonic relationship between fragment geometry and the required linker length. The remaining error, reflected by Relative Absolute Error (RAE) = 1.57, mainly arises from broad size variability among fragment pairs with similar anchor distances, where chemical topology and linker flexibility can lead to different numbers of linker atoms. These results show that the auxiliary modules provide useful geometric priors for FlexiTAC: the anchor module narrows the plausible attachment region, while the size module gives an informative estimate of linker length before generative sampling, allowing FlexiTAC to adapt to more scenarios when conditional information is lacking.

### S3 Supplementary Information for the PROTAC-DB 1.0 and ZINC evaluation results

To further examine whether FlexiTAC's performance arises from its algorithm strategy rather than from access to a larger in-house training set, we retrained and evaluated the model on two previously used benchmark datasets: ZINC and PROTAC-DB 1.0. In both settings, we followed the data splits, generation tasks and evaluation protocol used in the LinkerNet study, allowing a direct comparison under the same training and testing conditions.

Across both datasets, FlexiTAC remained highly competitive and often outperformed the original baselines. On ZINC (**Supplementary Table 1**), FlexiTAC achieved the highest validity and improved recovery over LinkerNet, while also showing lower median energy and RMSD after force-field relaxation. On PROTAC-DB 1.0 (**Supplementary Table 2**), FlexiTAC also improved recovery and maintained strong conformational quality under the same evaluation protocol. These results indicate that FlexiTAC's advantage is not simply due to the scale or curation of PROTAC-3D. Rather, its Bayesian flow network (BFN) formulation and structure-conditioned generation strategy provide a robust algorithmic benefit that transfers to datasets and configurations established by prior work.

**Supplementary Table-1| Evaluation Results on ZINC test set using the LinkerNet evaluation protocol.**

| Method | Valid, % | Unique, % | Novel, % | Rec, % | $E_{min}$ | RMSD | $E_L$ | $\Delta E_L$ |
|---|---|---|---|---|---|---|---|---|
| DeLinker | 96.8 ± 0.2 | 43.5 ± 0.4 | 43.3 ± 0.2 | 55.8 ± 1.2 | - | - | - | - |
| 3DLinker | 40.3 ± 0.1 | 53.6 ± 0.6 | 47.4 ± 2.7 | 43.2 ± 0.9* | ++ | 2.42 ± 0.01 | 5178.2 ± 108.5 | ++ |
| DiffLinker | 48.7 ± 0.1 | 90.1 ± 3.5 | 99.5 ± 0.0 | 0.0 ± 0.0 | 179.2 ± 6.7 | 1.92 ± 0.00 | 216.8 ± 1.0 | 93.3 ± 0.4 |
| LinkerNet | 83.1 ± 0.0 | 14.8 ± 0.1 | 11.4 ± 0.1 | 24.4 ± 0.1 | 32.7 ± 1.2 | 1.44 ± 0.00 | 54.8 ± 10.5 | 67.9 ± 1.3 |
| **FlexiTAC** | **98.9 ± 0.0** | 18.1 ± 0.0 | 60.8 ± 0.0 | 33.9 ± 0.9 | **23.6 ± 0.1** | **1.36 ± 0.00** | **52.6 ± 0.2** | **41.0 ± 0.1** |

**Supplementary Table-2| Evaluation Results on PROTAC-DB 1.0 using the LinkerNet evaluation protocol.**

| Method | Valid, % | Unique, % | Novel, % | Rec, % | $E_{min}$ | RMSD | $E_L$ | $\Delta E_L$ |
|---|---|---|---|---|---|---|---|---|
| DeLinker | 42.8 ± 0.4 | 89.9 ± 2.8 | **99.0 ± 0.1** | 0.0 ± 0.0 | - | - | - | - |
| DiffLinker | 24.0 ± 0.1 | **99.4 ± 0.0** | 98.9 ± 0.2 | 0.0 ± 0.0 | 416.2 ± 13.4 | 2.44 ± 0.05 | 501.0 ± 17.6 | 80.8 ± 4.6 |
| LinkerNet | 55.5 ± 0.0 | 47.9 ± 11.6 | 41.4 ± 4.9 | 5.4 ± 1.4 | 113.7 ± 8.0 | 1.55 ± 0.02 | **29.9 ± 17.1** | 500.0 ± 9.1 |
| **FlexiTAC** | **56.7 ± 0.1** | 51.5 ± 0.1 | 87.1 ± 0.2 | **23.3 ± 0.0** | **61.6 ± 12.3** | **1.37 ± 0.04** | 75.35 ± 23.0 | **44.61 ± 5.9** |

**Note**: Valid, Unique, and Novel denote standard 2D molecular generation metrics identical with **Fig. 2a**. Rec (Recovery rate) indicates the percentage of generated PROTACs that successfully recover the reference PROTACs. $\boldsymbol{E_{min}}$ represents the average minimum energy of generated molecules per fragment pair before RDKit MMFF optimization, indicating the overall quality of generated conformations. RMSD is the average Root Mean Square Deviation of molecule coordinates before and after MMFF optimization, reflecting the gap between generated conformations and the best possible ones. $\boldsymbol{E_L}$ denotes the average median energy of linker after MMFF constrained optimization with fragment atoms fixed. $\boldsymbol{\Delta E_L}$ represents the average median energy difference before and after this constrained optimization; a lower value indicates a better linker conformation. **As DeLinker is a 2D baseline model, 3D metrics are not applicable. All baselines and FlexiTAC were evaluated in triplicate, with ± denoting the standard deviation. The '++' symbol indicates exceptionally high energy values produced by 3DLinker.**

### S4 Supplementary Information for Jensen-Shannon divergence distributions of linker substructures

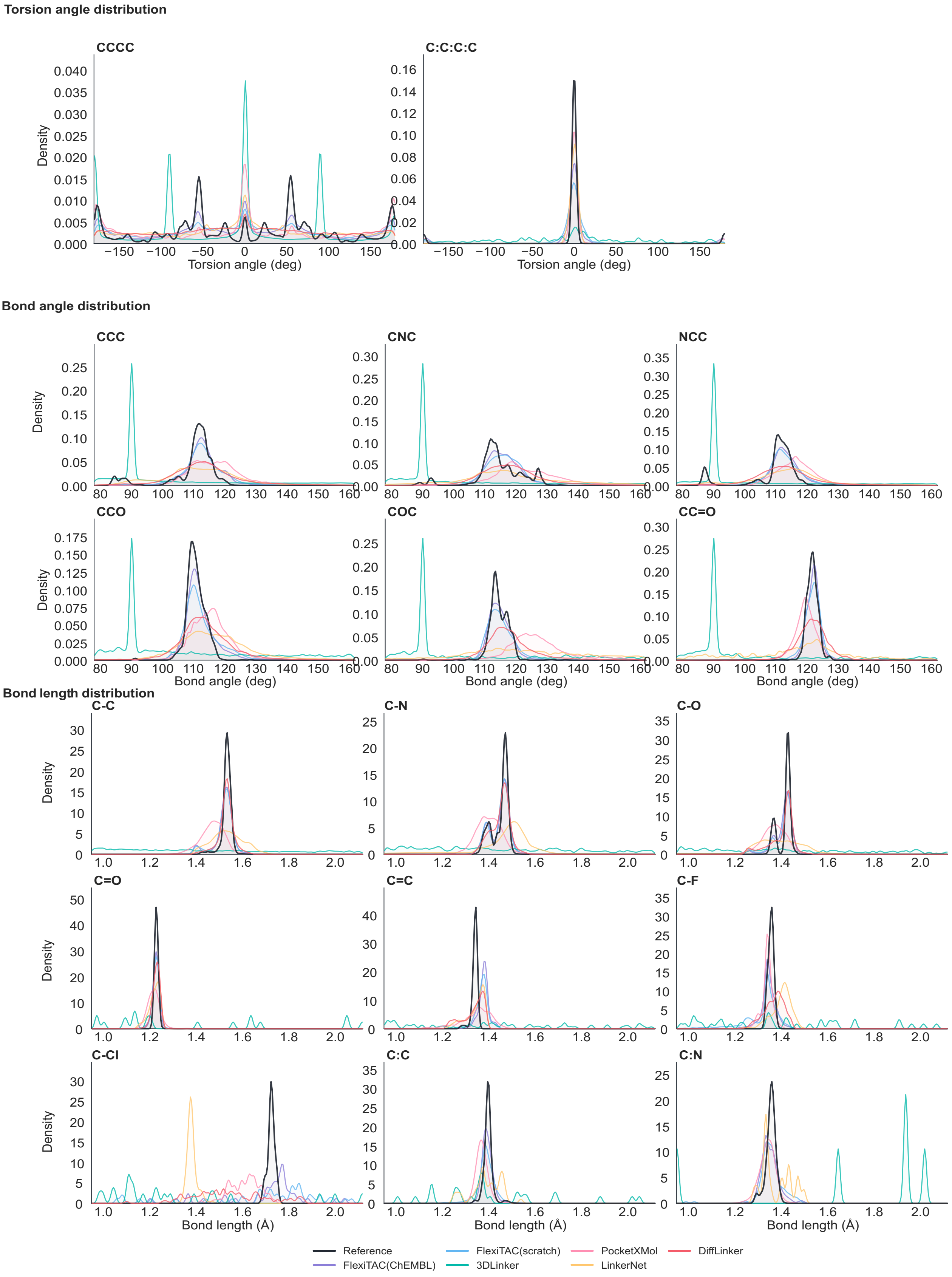

**Supplementary Fig. 6 |`Unsmoothed version of the Jensen-Shannon divergence distributions plot shown in the main text.** Density plots compare torsion-angle, bond-angle and bond-length distributions between reference linkers and molecules generated by different models. This is the none-smoothing version of the drifting figure in the main text.

### S5 Supplementary Information for motif coverage

**Reference linker motifs · Part 1**

**01. Piperazine**

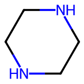


**02. Piperidine**

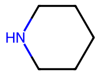


**03. Pyrrolidine**

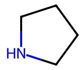


**04. Azetidine**

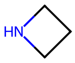


**05. Morpholine**

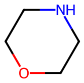


**06. 1,2,3-Triazole**

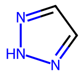


**07. 1,2,4-Triazole**

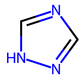


**08. Pyridine**

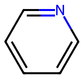


**09. Pyrimidine**

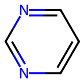


**10. Pyrazine**

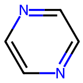


**11. Imidazole**

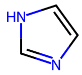


**12. Oxazole**

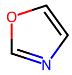


**13. Isoxazole**

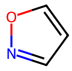


**14. Thiazole**

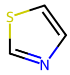


**15. Phenyl**

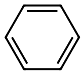

**16. Naphthalene**

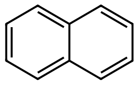

**17. Biphenyl**

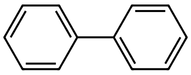

**18. Benzimidazole**

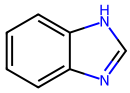


**19. Indole**

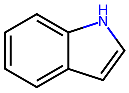


**20. Benzodioxole**

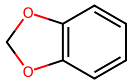


**21. Benzoxazole**

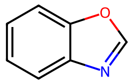


**22. Cyclobutyl**

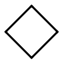

**23. Cyclopentyl**

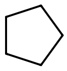

**24. Cyclohexyl**

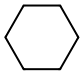

**25. Cyclopropyl**

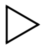

**26. Bicyclo[2.2.2]octane**

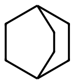

**27. 2,6-Diazaspiro[3.3]heptane**

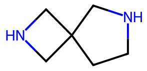


**28. 2,7-Diazaspiro[3.5]nonane**

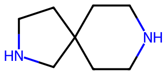


**29. Octahydro-pyrrolo[3,4-c]pyrrole**

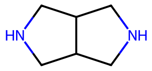


**30. Norborane**

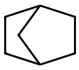

**31. PEG2**

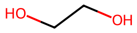

**Supplementary Fig. 7 | Motif structures from PROTAC-Bench test sets (Part 1 of 2)**. This figure presents the first 31 of 62 distinct motifs extracted to evaluate motif coverage on PROTAC-Bench. The complete set of 62 motifs reflects the diversity and rediscovery capabilities of linker design models at the substructural level.

### Reference linker motifs · Part 2

32. PEG3

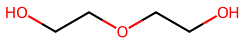


33. PEG4

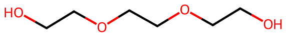


34. PEG5+

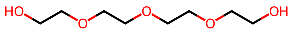


35. EG ether

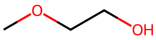


36. Amide

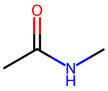


37. Urea

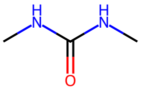


38. Sulfonamide

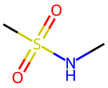


39. Ketone

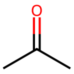


40. Ester

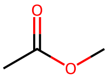


41. Ether

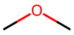


42. Thioether

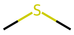


43. Acetylene

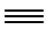

44. Propargyl ether

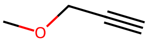


45. Nitrile

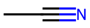


46. Fluoro




47. Chloro




48. CF3

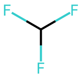


49. Hydroxyl




50. Primary amine

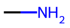


51. Test ring system RS01

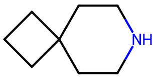


52. Test ring system RS02

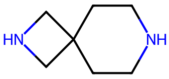


53. Test ring system RS03

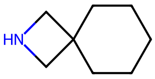


54. Test ring system RS04

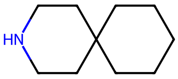


55. Test ring system RS05

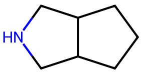


56. Test ring system RS06

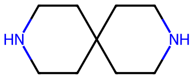


57. Test ring system RS07

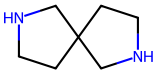


58. Test ring system RS08

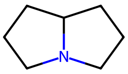


59. Test ring system RS10

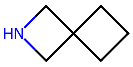


60. Test ring system RS11

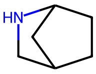


61. Test ring system RS12

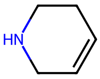


62. Test ring system RS13

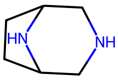

**Supplementary Fig. 8 | Motif structures from PROTAC-Bench test sets (Part 2 of 2). This figure presents the remaining 31 of 62 distinct motifs extracted to evaluate motif coverage on PROTAC-Bench. The complete set of 62 motifs reflects the diversity and rediscovery capabilities of linker design models at the substructural level.**

### S6 Supplementary Information for rigidity guidance sampling results

**Fragment Pair 1**

**Most Flexible**
γ = −1

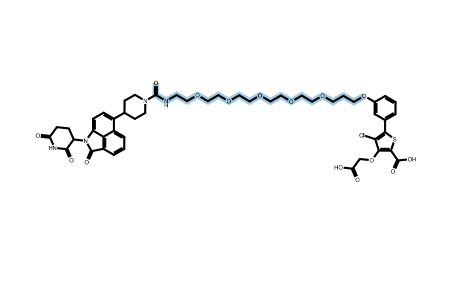

**Linker rigidity score: 0.2614**

**Ground Truth**
Reference

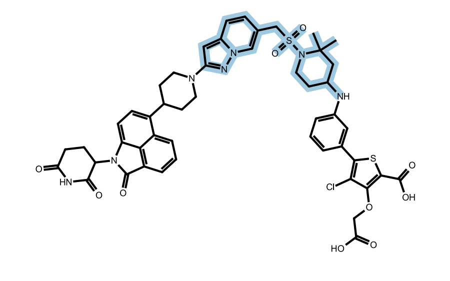

**Linker rigidity score: 0.4501**

**Most Rigid**
γ = +1

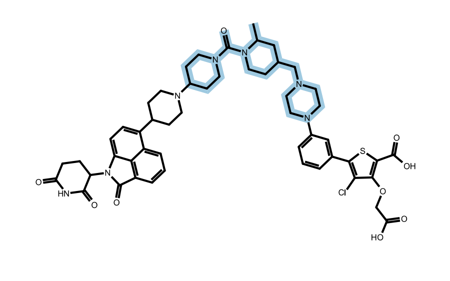

**Linker rigidity score: 0.8566**

**Fragment Pair 2**

**Most Flexible**
γ = −1

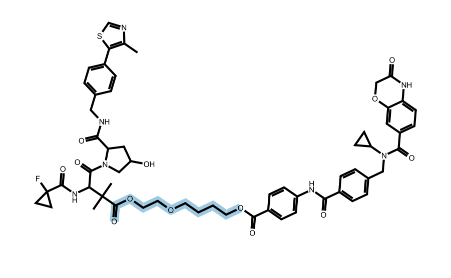

**Linker rigidity score: 0.4319**

**Ground Truth**
Reference

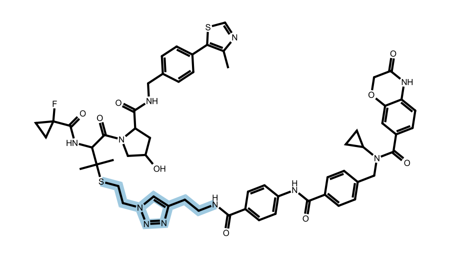

**Linker rigidity score: 0.6046**

**Most Rigid**
γ = +1

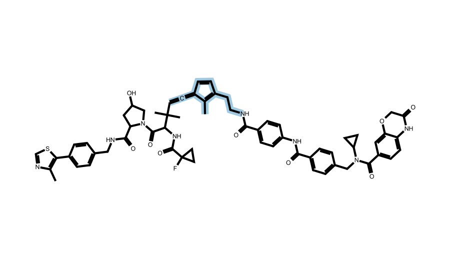

**Linker rigidity score: 0.8537**

**Fragment Pair 3**

**Most Flexible**
γ = −1

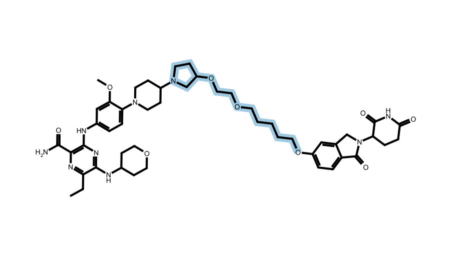

**Linker rigidity score: 0.3498**

**Ground Truth**
Reference

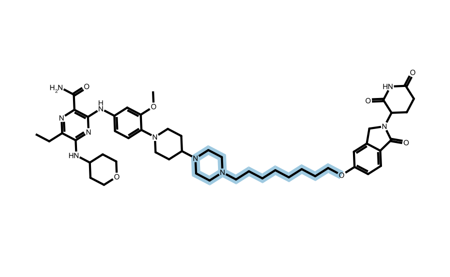

**Linker rigidity score: 0.4892**

**Most Rigid**
γ = +1

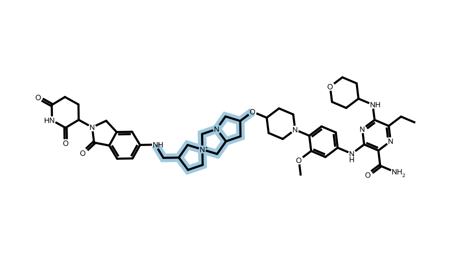

**Linker rigidity score: 0.8873**

**Fragment Pair 4**

**Flex-guided**
γ = -1.0

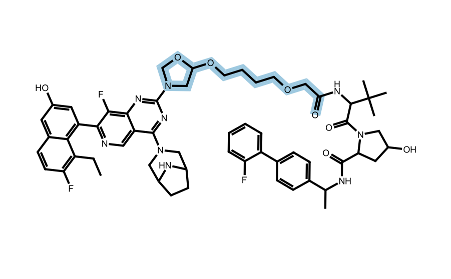

**Linker rigidity score: 0.5551**

**Ground Truth**
Reference

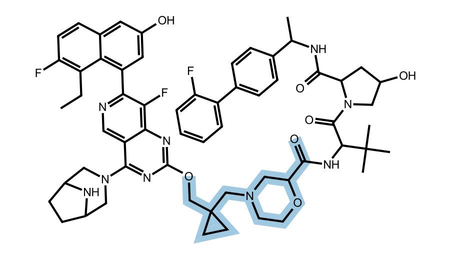

**Linker rigidity score: 0.6492**

**Rigid-guided**
γ = +1.0

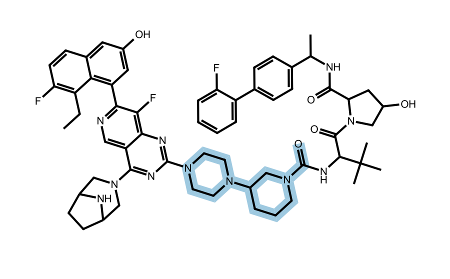

**Linker rigidity score: 0.8156**

**Supplementary Fig. 9** | Representative PROTACs generated under different rigidity-guidance. For each fragment pair, FlexiTAC was sampled with flexible guidance ($\boldsymbol{\gamma} = -\mathbf{1}$) and rigid guidance ($\boldsymbol{\gamma} = +\mathbf{1}$), and the generated PROTACs were compared with the corresponding ground-truth molecule. The flexible-guided samples generally contain more extended and rotatable linker regions, whereas the rigid-guided samples show higher rigidity scores and more constrained linker architectures. These examples illustrate that rigidity guidance can shift linker flexibility while preserving the same fragment-pair context.

**Fragment Pair 5**

**Most Flexible**

γ = −1

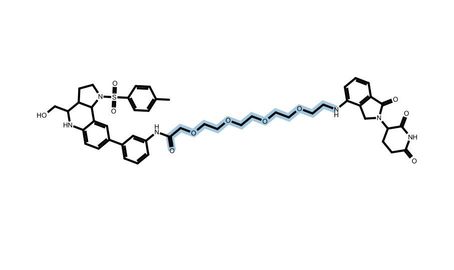

**Linker rigidity score: 0.3175**

**Ground Truth**

Reference

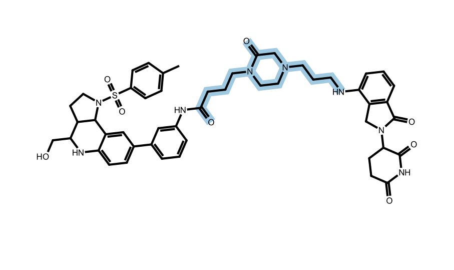

**Linker rigidity score: 0.5548**

**Most Rigid**

γ = +1

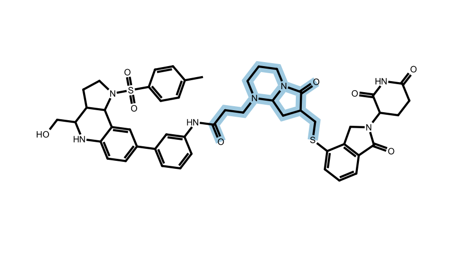

**Linker rigidity score: 0.8003**

**Fragment Pair 6**

**Most Flexible**

γ = −1

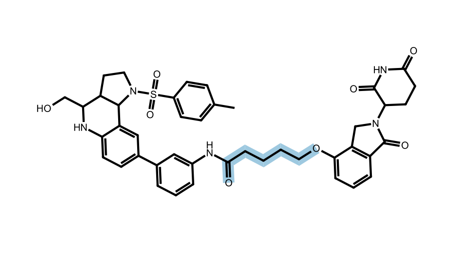

**Linker rigidity score: 0.5788**

**Ground Truth**

Reference

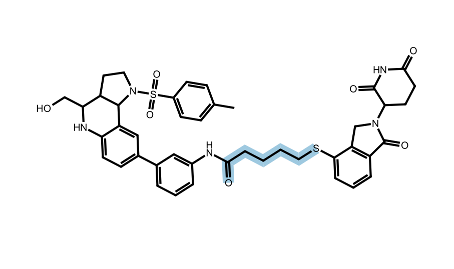

**Linker rigidity score: 0.6300**

**Most Rigid**

γ = +1

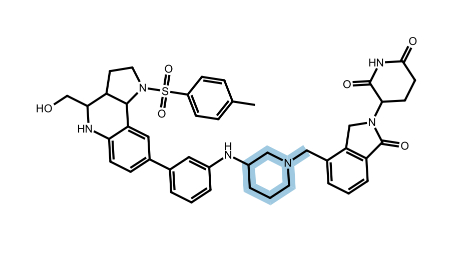

**Linker rigidity score: 0.9994**

**Fragment Pair 7**

**Most Flexible**

γ = −1

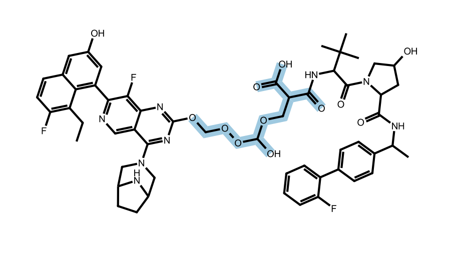

**Linker rigidity score: 0.3976**

**Ground Truth**

Reference

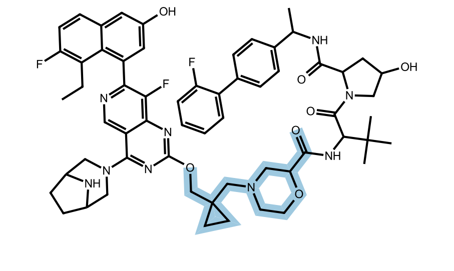

**Linker rigidity score: 0.6492**

**Most Rigid**

γ = +1

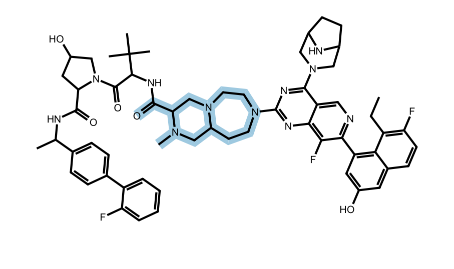

**Linker rigidity score: 0.9292**

**Fragment Pair 8**

**Flex-guided**

γ = -0.5

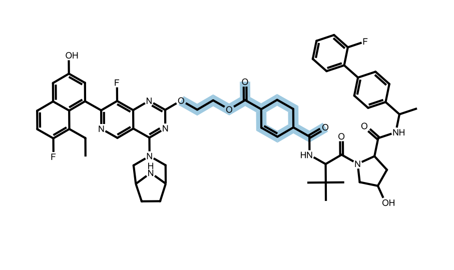

**Linker rigidity score: 0.6491**

**Ground Truth**

Reference

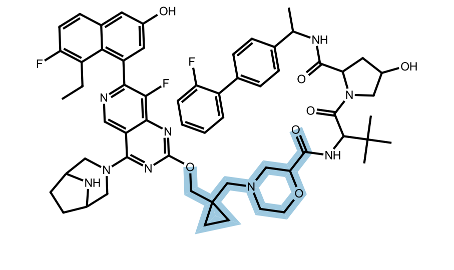

**Linker rigidity score: 0.6492**

**Rigid-guided**

γ = +1.0

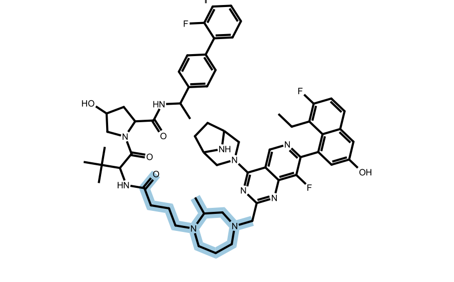

**Linker rigidity score: 0.6683**

Supplementary Fig. 10 | Representative PROTACs generated under different rigidity-guidance settings (Part 2 of 2). Additional examples of fragment pairs are presented. FlexiTAC was sampled with flexible guidance ($\boldsymbol{\gamma} = -\mathbf{1}$) and rigid guidance ($\boldsymbol{\gamma} = +\mathbf{1}$), and the generated PROTACs were compared with the corresponding ground-truth molecule. These examples further illustrate that rigidity guidance can effectively shift linker flexibility, yielding more extended or constrained architectures, while preserving the same fragment-pair context.

### S7 Supplementary Information for the multi-scenario application of FlexiTAC

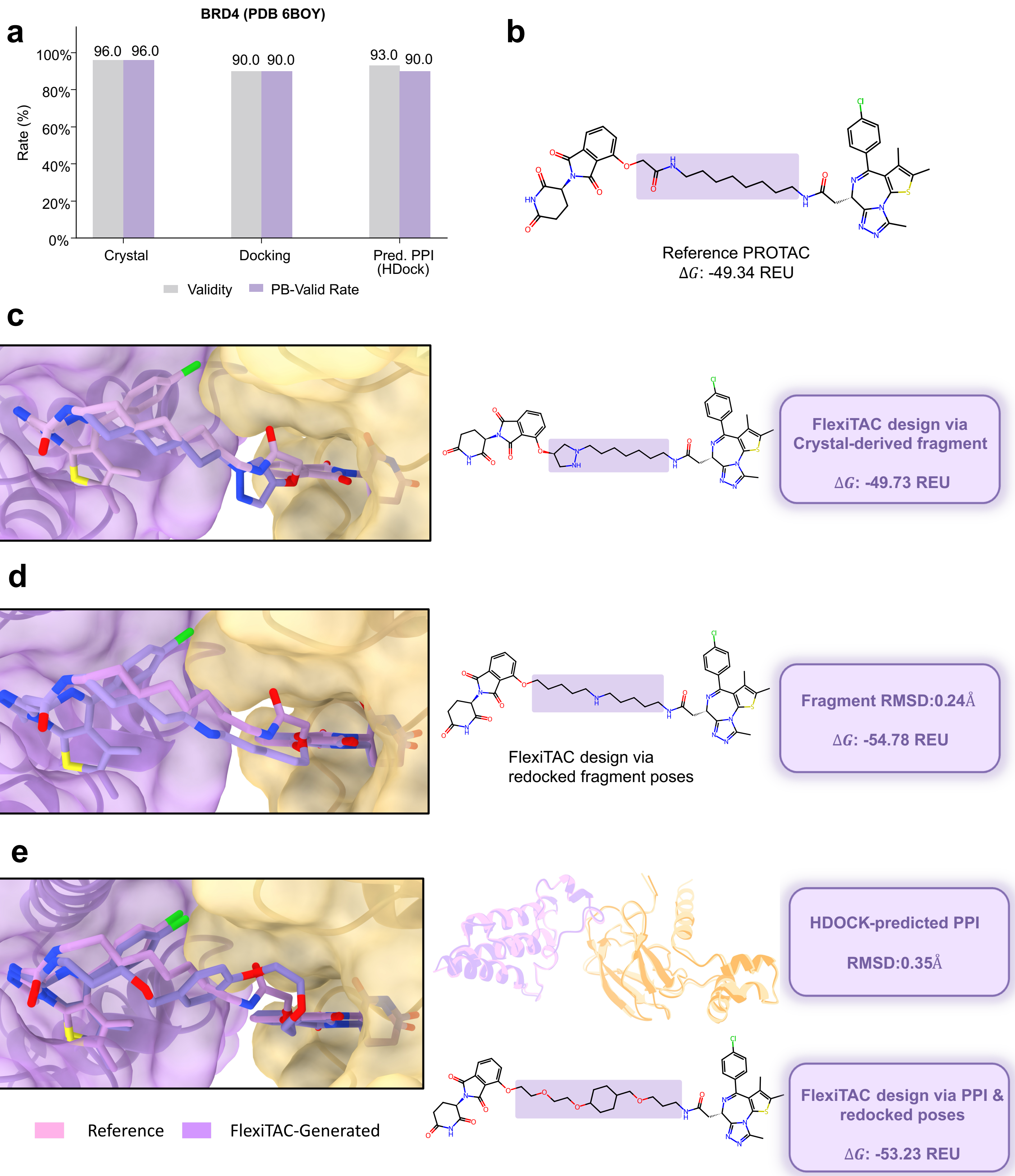

**Supplementary Fig. 11** | FlexiTAC linker design for BRD4 (PDB 6BOY) using multiple structural inputs. **a,** Chemical validity (grey) and PoseBusters pass rates (purple) for designs generated from crystal, redocked and HDock-predicted inputs. **b,** Reference PROTAC from the 6BOY crystal structure, with the linker highlighted (ΔG, -49.34 Rosetta Energy Unit (REU). **c,** FlexiTAC design generated via crystal-derived fragment poses (ΔG, -49.73 REU). **d,** FlexiTAC design generated via redocked fragment poses (fragment RMSD, 0.24 Å; generated ΔG, -54.78 REU). **e,** FlexiTAC design via HDOCK-predicted protein-protein interaction (PPI) structure and redocked poses (PPI RMSD, 0.35 Å; generated ΔG, -53.23 REU). RMSD were calculated between the crystal and HDOCK-predicted PPI structure, and ΔG values were calculated using PRosettaC. Reference and FlexiTAC-generated PROTACs are shown in pink and purple, respectively. All three generated designs had more favourable calculated ΔG values than the crystal reference.

### S8 Supplementary Information for baseline models on PROTAC-Bench

PROTAC-Bench contains seven baseline models that are grouped into two main categories. The 2D baselines are DeLinker and Link-INVENT. DeLinker is a graph-based linker-generation model that operates on molecular topology and uses the fragment and their attachment geometry as conditions. Link-INVENT is a sequence-based linker generator built within the REINVENT framework and was evaluated through the REINVENT4 sampling pipeline. The 3D baselines are 3DLinker, DiffLinker, LinkerNet, FFLOM and PocketXMol. 3DLinker was treated as a linker-specific 3D baseline because it explicitly generates linker topology and 3D coordinates from fragment conformations. DiffLinker and LinkerNet were equivariant diffusion baselines. PocketXMol is a pretrained 3D molecular generation framework adapted to the PROTAC linker setting through a constrained sampling strategy. Among all the baselines, DeLinker, Link-INVENT, 3DLinker, DiffLinker and LinkerNet were reported as models specifically designed for linker-generation task, while FFLOM and PocketXMol were considered as versatile models that capable of linker-design tasks.

Regarding the training and inference settings, five baselines were retrained on the PROTAC-3D training data the same as FlexiTAC(scratch): DeLinker, 3DLinker, DiffLinker, LinkerNet and FFLOM. All retraining used the original model implementations, losses and recommended training workflows. Link-INVENT and PocketXMol were not retrained. Link-INVENT was sampled from the released pretrained checkpoint used by the REINVENT4 workflow using reinforcement learning, and PocketXMol was sampled from its released checkpoint pretrained on large-scale data containing small molecules, peptides and proteins. For evaluation, all methods generated 100 linkers per fragment pair, leading to a total number of 13,000 PROTACs for benchmark evaluation. Generation was performed on the three PROTAC-3D test settings: both-unseen, E3-unseen and warhead-unseen. The same fragment-pair definitions and reference PROTAC sets were used for all models.

During generation, reference linker size is hard-coded or explicitly supplied in all baseline sampling pipelines. Anchor-atom labels are also required to specify the two attachment atoms connecting the generated linker to the input fragments. To ensure a controlled comparison across heterogeneous model interfaces, we supplied the same reference linker size and anchor-atom information to baseline models.

For the model-specific implementations, DeLinker was retrained on the PROTAC-3D train set using its graph-generation objective, while 3DLinker was retrained with the PROTAC-3D training configuration and evaluated as the linker-specific 3D baseline. Both DiffLinker and LinkerNet were retrained with the PROTAC-3D anchor-aware configuration, with LinkerNet specifically evaluated as an equivariant diffusion model for fragment-pose and linker co-design. FFLOM was retrained using processed PROTAC-3D tensors and molecule SMILES, and its sampling setup fixed the linker-length freedom to match the reference linker-size condition. Link-INVENT and PocketXMol were evaluated directly from their released pretrained checkpoints.
